# Characteristic core voxels in normal individuals revealed by hyperbolic disc embedding and *k*-core percolation on resting state fMRI

**DOI:** 10.1101/2021.08.15.456381

**Authors:** Wonseok Whi, Youngmin Huh, Seunggyun Ha, Hyekyoung Lee, Hyejin Kang, Dong Soo Lee

**Affiliations:** Department of Molecular Medicine and Biopharmaceutical Sciences, Seoul National University; Department of Nuclear Medicine, Seoul National University and Seoul National University Hospital; Medical Research Center, Seoul National University; Division of Nuclear Medicine, Department of Radiology, Seoul St. Mary’s Hospital, College of Medicine, The Catholic University of Korea; Biomedical Research Institute, Seoul National University Hospital

**Keywords:** Resting-state fMRI, Independent components-voxels composition, *k*-core percolation, *k*_max_-core, Hyperbolic disc embedding

## Abstract

Hyperbolic disc embedding and *k*-core percolation reveal the core structure of the functional connectivity on resting-state fMRI (rsfMRI). Inter-voxel relations were visualized on embedded hyperbolic discs, and their core composition was traced using *k*-core percolation. Using 180 normal adults’ rsfMRI data from the Human Connectome Project database, scale-free intervoxel connectivity represented by IC-voxels composition, while visualized on hyperbolic discs using 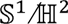 model, showed the expected change of the largest component decreasing its size on *k-*core percolation eventually yielding the core structures of individuals. This *k*_max_-core voxels-ICs composition revealed such stereotypes of individuals as visual network dominant, default mode network dominant, and distributed patterns. Characteristic core structures of resting-state brain connectivity of normal subjects disclosed the distributed or asymmetric contribution of voxels to the *k*_max_-core, which suggests the hierarchical dominance of certain IC subnetworks characteristic to subgroups of individuals at rest.

## Introduction

Brain composes a high-dimensional complex and integrated network, which is composed of multiple modular and specialized networks, distributed spatially, and combined to form a multi-modular structure [1–5]. The conundrum of how these modules are aggregated to form a single coherent network with preserved functionality remains a fundamental question for unveiling the functional architecture of the human brain at resting-state and on activation [6, 7]. Individual differences add complexity to a succinct understanding of this question.

Recent works of network science suggest that the aggregation of these modules is facilitated by a set of essential voxels, which integrate intra-modular and inter-modular information throughout the network [2, 3, 8, 9]. Essential nodes were at first supposed to be the hub nodes with high degrees on the brain graphs but soon were suggested to be rather the core influencer nodes with a wide range of initial degrees on decomposition [8]. In physical networks, disruption in these core nodes led to abrupt disintegration having been called network dismantling or targeted damage [10–13]. The disruption in the brain graph is associated with serious neuropsychiatric diseases with disrupted associative functionality [14–15].

Therefore, it is important to identify which nodes compose the core structure, and if ever the resting state core is individually unique, then their individual differences of the core composition should be disclosed using voxel-based representation of brain graphs. Recent studies of mathematics and neurosciences have accomplished this job successfully with physical networks and probably with the brain [3, 8, 12–16], implementing hubness, centrality measures such as degree, betweenness, eigenvector, and leverage centrality [17–20], or *k*-core [21–25]. The *k*-core percolation describes the architecture of the backbones of the network, by filtering out peripheral nodes and searching for remaining central nodes, where the coreness, *k*, acts as the threshold for sustaining node connectivity along the filtration. The *k*-core percolation was used to understand the forward (phase transition) and backward (k- decomposition) behavior of networks and brain networks can be dissected in a similar way as was done for graph filtration thresholding [26–30].

In our previous work, we tackled the problem of difficulty in visualization and thus forming mental imagery of the object brain graphs by considering their geometric characteristics. We adopted an analytical framework for visualizing a complex, multi-modular network for functional brain networks with scale-freeness, by embedding the networks into the latent geometric model of 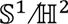 representing them on a hyperbolic space [31–33]. This solved two issues, 1) 2-dimensional representation of complex brain graphs with flexible annotation of functionally cooperating voxel groups and 2) thresholding to sort out necessary edges to make the complex brain graphs obey power law and thus scale-freeness [34]. Map of functional brain network acquired by resting-state functional magnetic resonance imaging (rsfMRI) was visualized using hyperbolic disc model, and we could estimate functional proximity/homogeneity between voxels by a measure of hyperbolic distance on this disc [31–34].

In this work, we analyzed functional brain networks from healthy human young adults by analyzing rsfMRI data and visualized functional subnetworks, i.e., independent components (ICs), using the hyperbolic embedding. Then, we investigated how each functional subnetwork was composed of the subset of voxels with high core-ness, revealed by *k*-core percolation as a measure of centrality. We characterized the *k*_max_-core voxels for their degree distribution belonging to each IC, showing the plausible influencer behavior of these IC subnetwork voxels upon *k*-core percolation eventually to find which subnetworks are the dominant by counting the voxels belonging to them at rest in normal individuals. We asked whether the individuals had common or characteristic core structures in terms of their *k*_max_- core IC-voxels compositions.

## Results

### Method of hyperbolic embedding of voxels on individual rsfMRI

To visualize the correlation structure of the voxels composition of the complex functional brain network, we adopted a method to transfer the high-dimensional connection (edge) information to the hyperbolic disc space. According to our previous investigation [34] having looked for an optimal non-Euclidean space for embedding the inter-voxel correlation structure, we simply chose 2-dimensional hyperbolic disc embedding. Hyperbolic disc representation reflected the original high-dimensional edge information similarly well to the high dimensional Euclidean embedding alternatives [34]. Hyperbolic disc embedding rendered a high-dimensional correlation structure to have the easy-to-recognize visibility, which had never been achieved using Euclidean 2-dimensional representation (Fig. 1a, b).

**Figure 1.**
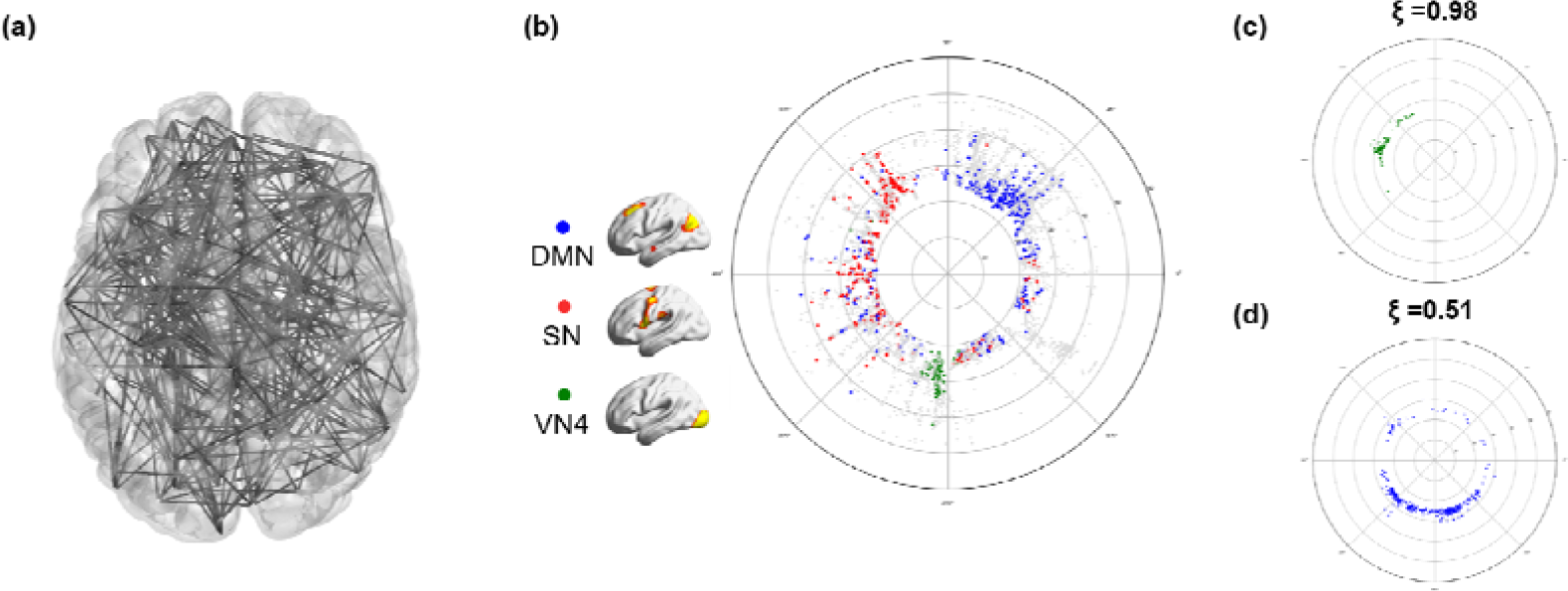
Hyperbolic disc embedding and angular coherence of the voxels on the disc. (a) A brain network with random 500 nodes was displayed using 3-dimensional MNI coordinates projected on a 2-dimensional brain space. This visualization provides intricate edges and nodes not easily discernable. (b) Hyperbolic disc embedding provides easy-to-recognize visualization of the voxels on the hyperbolic disc. In this hyperbolic disc, 5,937 voxels were used, which shows the inter-voxel relationships between voxels in an un-overlapped way with 10 times more voxels than the one in (a). Specifically, the brain was resampled into 5,937 6 × 6 × 6m voxels, which were assigned as voxels of subnetworks belonging to independent components (ICs) [34]. The hyperbolic distance between two voxels on this hyperbolic disc is equivalent to the correlation proximity between these voxels on the Euclidean space. The radial coordinate responds to the degree of the voxel, i.e., the hub voxel is near to the center of the disc, and the angular coordinate responds to the similarity of voxels [32]. As an example, voxels from the independent component (IC) subnetworks were presented in different colors. The voxels from the salience network (SN) (red) show more widely distributed on angular coordinates than the default mode network (DMN) (blue) or visual network 4 (VN4) (green). (c) The angular coherence quantifies the degree of aggregation of a group of voxels based on coordinates. It ranges from 0 to 1, and a higher value indicates compact gathering with smaller differences in angles between voxels in the group. Widely spread voxels have a lower value of angular coherence. The angular coherences of voxels comprising (c) VN4 ( =0.98) (c) and (d) DMN ( =0.51) are shown.

Unlike our previous study which used anatomically predefined regions [34], we used voxels’correlation to visualize inter-voxel relationship for hyperbolic disc embedding using 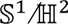 model [31]. The output was easy to disclose the belonging characteristics of the voxels to ICs on a hyperbolic polar coordinate. Edge weight on the voxel-voxel correlation matrix was thresholded to yield the adjacency matrix after confirming the scale-freeness of the resultant degree distribution of the voxels, while preserving the size of the largest component as large. This allowed us to confirm the power-law of its degree distribution and scale-freeness to be fit for the hyperbolic model. Reproducibility on repeated embedding was tested in an exemplary case with repeated embedding with Mercator algorithm [31, 34] (Supplementary Figure 1).

### Hyperbolic disc embedding of rsfMRI voxels and their belonging to IC subnetworks

We downloaded the resting-state fMRI data of 180 adults without any psychiatric or neurological diseases from the Human Connectome Project (http://www.humanconnectomeproject.org/) and produced inter-voxel correlation networks with 5,937 voxels. Raw voxels were down-sampled to yield computationally plausible size but we still called the enlarged units of aggregates of voxels as voxels. The correlation coefficients between two voxels were calculated to define the edges of the network. These networks were binarized after confirming the linearity on a log-log plot of the degree distribution of the output adjacency matrix, and the largest components of the network of having at least 80% of the entire 5,937 voxels were embedded on the hyperbolic discs using the previously described method [34]. Embedding was done on the hyperbolic disc using 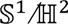 model according to the methods previously reported [31, 34].

BOLD time series of voxel pairs were used to calculate inter-voxel correlation and after confirming scale-freeness on the degree distribution, the correlation matrix was thresholded to yield the adjacency matrix. This adjacent matrix was then converted to fit into 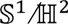 model finally to yield the polar coordinates for the voxel. Using the polar coordinates, we estimated angular coherence to investigate how the embedded voxels were angularly similar. After successful embedding using Mercator [31], voxels included in the largest components were more than 80% (5,391 ± 224 voxels) with the edges ranged from 274,634 to 3,894,033 (1.6 ∼ 22% of possible edges). A randomly sampled case was repeatedly embedded to this 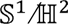model and their reproducibility was shown in Supplementary Figure 1 and Supplementary Table 1. Voxel-based embedding in this study yielded a similar feature of reproducibility. We confirmed that 2-dimensional hyperbolic disc embedding using 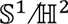 model was feasible using voxel-based data and analyzed the pattern of embedded voxel-voxel relationships. We used only the positive correlation and the negative correlation were left for the following study, including also the inter-dependent multi-layered characteristics of brain networks on hyperbolic disc embedding.

This embedding provided clearer visibility of inter-voxel relations on 2-dimensional space (disc) than any conventional method of visualization (Fig. 1a, b). Initial suggestion of using hyperbolic disc for popularity and similarity representation of growing complex network and translation of this method to 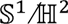model [32] enabled us to assume that the angular coherence on the hyperbolic disc reveals similarity of the group of voxels and that the distance of a voxel to the disc center represents higher degree with greater popularity [34]. We identified the voxels belonging to specific ICs (fifteen ICs) obtained from conventional group ICA performed in the entire 180 subjects [36].

### Angular coherence of hyperbolic disc-embedded voxels belonging to IC subnetworks

We investigated the distribution pattern of voxels on the hyperbolic discs and their belongings to each IC among normal individual subjects. Group ICA annotated each voxel to its IC. Angular coherence of grouped voxels according to ICs was measured on embedded hyperbolic discs. When voxels belonging to an IC were grouped closely together within narrow angles from the disc center, the IC and its voxels were called to have higher proximity with higher angular coherence (ranging from 0 to 1) (Fig. 1c, d). Angular coherence of ICs represented how close the voxels in an IC gathered together as a subnetwork. Subnetworks were labeled in two ways, one with the functional label of the voxels to the 15 ICs reminded the multiscale renormalization of brain graphs [39] (Supplementary Figure 2). Another anatomical label used predefined 15 lobes based on the Brainnetome atlas [40] (Supplementary Figure 3).

For the 180 subjects (Supplementary Table 2), the angular coherence calculated using functional labels tended to be higher than that using anatomical labels. In the functional label, sensorimotor network 1 (SMN1) (∼0.81), VN3 (∼0.80)/5 (∼0.75) showed the highest angular coherence, and precuneus network (PCN) (∼0.48) and dorsal attention network (DAN) (∼0.50) showed the lowest (Fig. 2a, Supplementary Table 3). Bilateral occipital lobes (left: ∼0.71, right: ∼0.72) showed the highest angular coherence, and parietal (left: ∼0.25, right: ∼0.26) and temporal lobes (left: ∼0.29, right: ∼0.25) showed the lowest in the anatomical label (Fig. 2b, Supplementary Table 4). The number of voxels belonging to the 15 ICs ranged from 158 (anterior default mode network (aDMN)) to 443 (visual network (VN) 2) (Supplementary Table 3). On the hyperbolic disc, angular coherence (ξ. of DMN were 0.57 ± 0.14 (n=180) and that of VN1 was 0.71 ± 0.15. In an individual chosen, for example, VN4 (green circle) voxels had ξ = 0.95, salience network (SN, red circle) ξ = 0.52 or the default mode network (DMN, blue circle) ξ = 0.50. The voxels embedded on the hyperbolic disc disclosed their own unique pattern but also revealed the common distribution characteristics. The hyperbolic disc should be read with polar coordinates with its inter-voxel distance on a logarithmic scale in the radial direction and hyperbolic contribution of inter-voxel angle in arc hypercosine to the distance (See Methods). Evidently, there should be considered the rotation/reflection symmetry [37] of this embedded disc representation and also other symmetry such as branch permutation related with the hyperbolicity of disc in the interpretation of voxels distribution [38].

**Figure 2.**
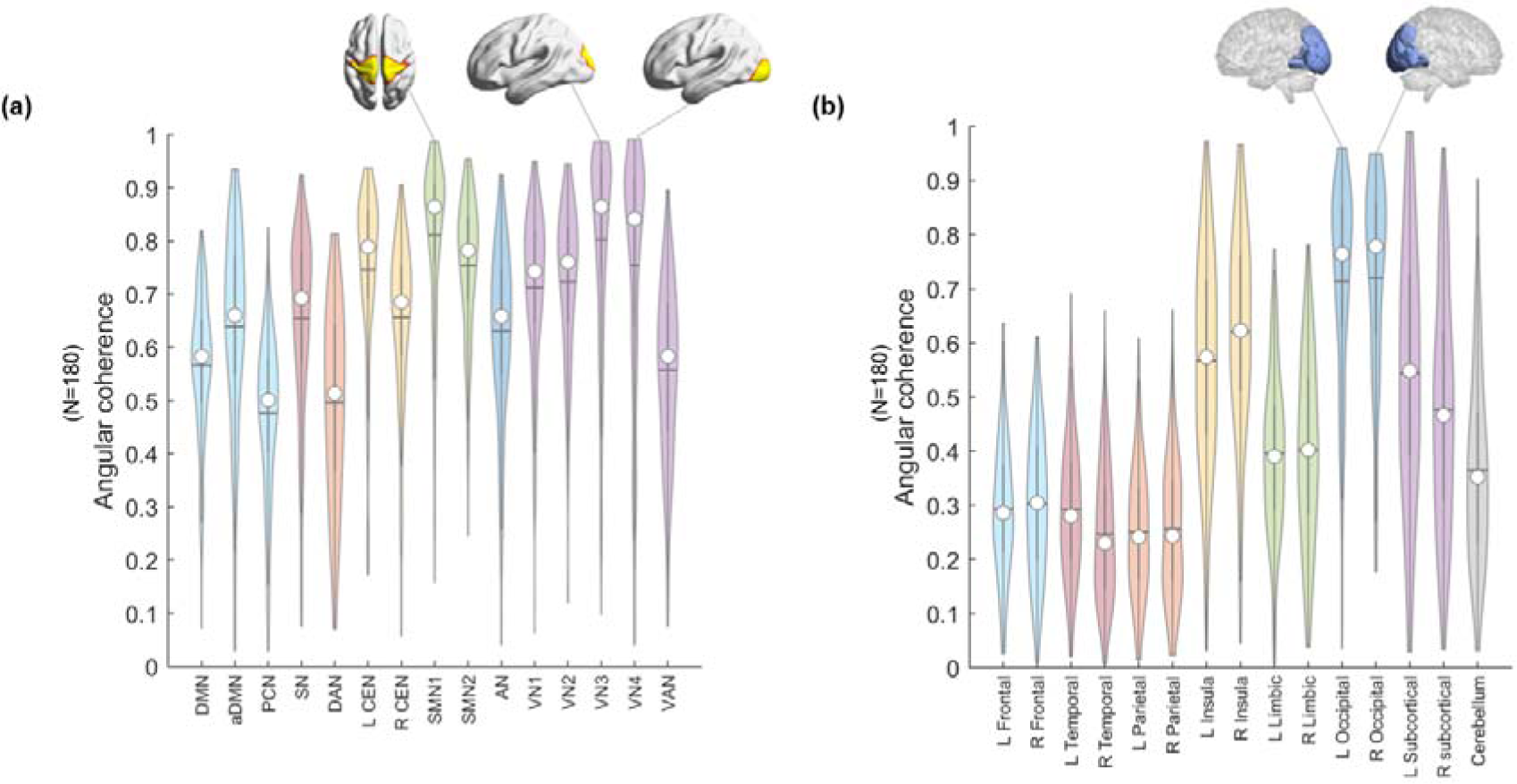
Distribution of the angular coherences of 180 individuals’ voxels on hyperbolically embedded discs. Groups of voxels belonged to (a) functional labels derived from group independent component analysis (ICA) and (b) atlas-based anatomical labels. (a) Fifteen independent components were chosen from group ICA for the entire data, which were Default mode network (DMN), anterior DMN (aDMN), precuneus network (PCN), salience network (SN), dorsal attention network (DAN), left central executive network (L CEN), right CEN (R CEN), sensorimotor network (SMN) 1/2, auditory network (AN), visual network (VN) 1/2/3/4, and visual attention network (VAN). The spatial maps of ICs were binarized (Z > 6), and voxels were classified to belong to each of the specific ICs. The coordinates of groups of voxels per specific ICs were calculated (see Methods). The values of angular coherence of SMN and VN3, VN4 was the highest. (b) Whole-brain was segmented into fifteen anatomical lobes based on Brainnetome atlas to yield anatomical labels: bilateral frontal/temporal/parietal/limbic/occipital/subcortical and a cerebellum. The coordinates of groups of voxels per a certain lobe were calculated (see Methods). The median of each distribution was indicated with a circle and the mean with a horizontal line.

Regarding laterality, the angular coherence of each left and right hemisphere based on the Brainnetome atlas, except cerebellum, showed no significant difference (Supplementary Figure 4a). An analysis using the anatomical label, temporal lobes, and subcortical regions showed significantly different angular coherence between the left and the right (*p* < 0.05, Supplementary Figure 4b).

### Discovery of k_max_-core voxels on k-core percolation and its visualization on hyperbolic embedded discs

Using the adjacency matrix obtained by scale-freeness guaranteed thresholding, we proceeded to find the core voxels. We asked which voxels in the IC subnetworks survived the decomposition by *k*-core percolation. We questioned whether the hub voxels with higher degrees would remain solely or other voxels with fewer edges join the survivors. And which voxels belonging to which ICs would remain and dominate or participate as core voxels at the end of *k*-core percolation.

During *k*-core percolation, voxels with a degree *k* were designated as *k*-shell and removed, and *k* started from 1 with an increment of 1. This *k-*shell removal was accompanied by recalculation of the remaining voxels’ degrees and was repeated until the step of *k_max_*; at this maximum step when one step further, there would have been left no largest component. This procedure peels the layers of a network based on the n-degree of voxels. The voxels with a degree equal to the coreness *k* are called *k*-core, and voxels with the highest degree at the step of coreness *k*_max_ are called *k*_max_-core (Fig. 3a, c). The *k*_max_-core voxels were not always the voxels with the highest degree at the beginning (Fig. 3b) since *k*-percolation sequentially eliminated voxels with lower degrees than the *k* and recalculation made voxels survive or not with their remaining connections with then-survivors (Fig. 4).

**Figure 3.**
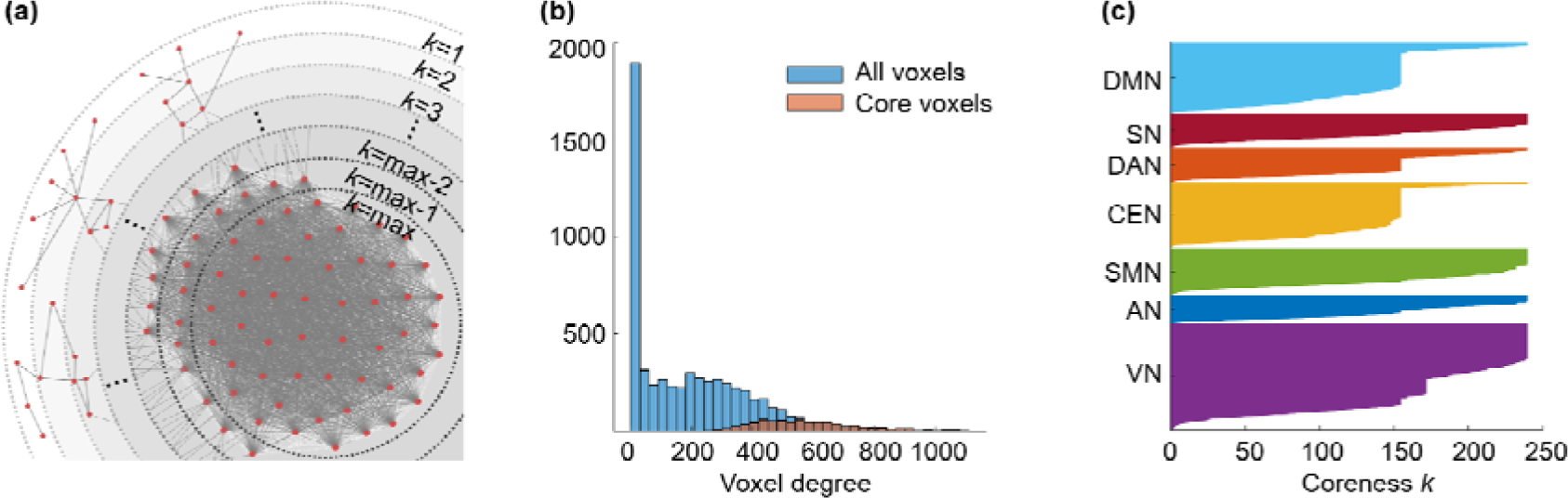
Conceptual illustration of *k*-core percolation and plots describing *k*-cores and the *k*_max_-core derived by *k*-core percolation. (a) *k*-core percolation renormalizes the brain network by peeling the layers with *k*-step from *k*=1 to *k*=max for the brain network. Intervoxel correlations were thresholded to yield an adjacency matrix after checking the scale-freeness of the degree distribution of voxels and put into hyperbolic disc embedding and *k*-core percolation. The voxels with a degree equal to coreness *k* are eliminated and recalculation of the voxels’ degree proceeded to the next step and went on until the remaining voxels forming the largest component at that step came to be disintegrated into many pieces at once. The voxels at this step *k*=max are called *k*_max_-core voxels. (b) The *k*_max_-core voxels included not only the voxels with the largest degree on the initial adjacency matrix but also the voxels with smaller degrees. This histogram shows the degree distribution of voxels from one subject (#100206). The blue bins represent all the voxels and the red for *k*_max_-core voxels. *k*_max_ was 240 and the degrees of *k*_max_-core voxels ranged from 260 to 1088. The *k*-core percolation finds *k*_max_-core voxels that have dense connectivity among themselves as well as hierarchically at the apex within their belonging independent components (ICs) and even the voxels with lower down to one-fourth of voxel with the highest degree. (c) A flags-plot shows the changing *k*-cores of a subject that vary with coreness *k* value during *k*-core percolation. Each voxel that belongs to a specific IC was shown on the y-axis, and the voxels comprising each *k*-core were colored. This subject has a *k*_max_-core with a 240 *k*-value and shows the first abrupt decrease during *k*-core percolation in DMN, DAN, CEN, and VN (*k* 156), and the second abrupt decrease in VN (*k* 172).

**Figure 4.**
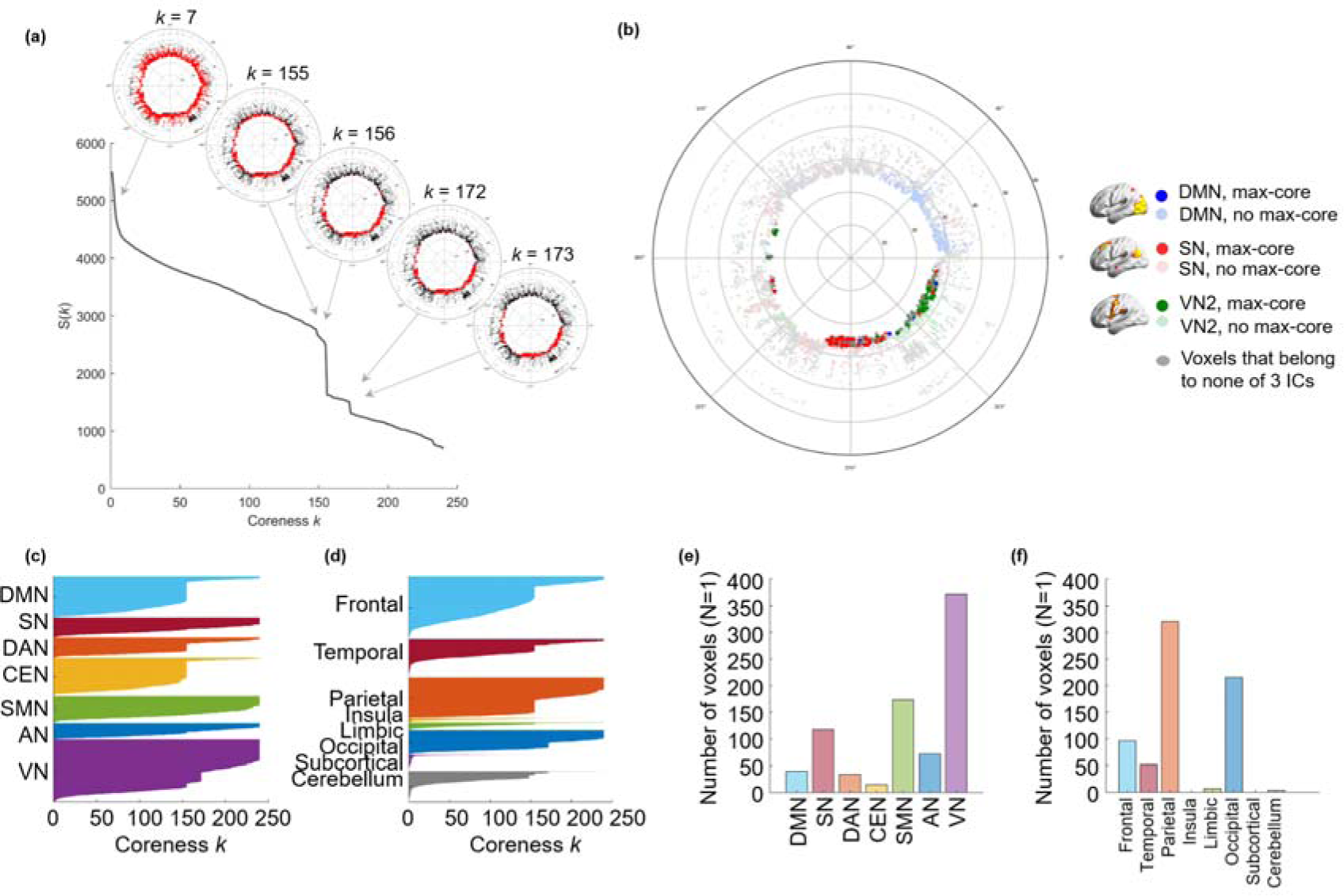
The *k*-cores and the *k*_max_-core depicted by flags-plots and hyperbolically embedded discs. Each individual has his/her own size of *k*_max_-core and changes in the size of *k*-cores according to coreness *k* during *k*-core percolation. In individuals, a few abrupt decreases were observed over the gradual change of the largest component. (a) The coreness *k* and the size of the core S(*k*) of a subject were plotted showing two abrupt changes. Specific *k*-cores that showed an abrupt decline (*k*=7, 155, 156, 172, 173) were embedded on the hyperbolic discs to show the explosive decrease of core voxels. The voxels belonging to a *k*- core were denoted with red circles, otherwise with black circles, on these hyperbolic discs. When the plot shows an abrupt decrease of S(*k*), voxels belonging to the *k*-core are reduced at once. (b) In an individual, *k*_max_-core shows the various size and the independent component (IC)-voxels composition. The *k*_max_-core (*k*=240) of a subject was presented as an example. There were 694 voxels left on the *k*_max_-core, and the voxels that belong to the default mode network (DMN) were in blue, salience network (SN) in red, visual network 3 (VN3) in green. Other voxels than *k*_max_-core voxels were in pale circles. (c) The components of each *k*-core from one subject that vary with coreness *k* value were shown on the flags-plot using functional IC labels (c) and anatomical labels (d) that annotated voxels to specific subnetworks. In the flags-plot, every voxel is presented in y-axis with labeling, and the horizontal bar of each voxel refers to the maximum *k* of *k*-cores that the voxels belong to. The voxels from each subnetwork on y-Axis were sorted in descending order of *k*. The bar plots show the affiliation of *k*_max_-core voxels (e, f). This individual showed abrupt declines of *k*- core size in DMN, dorsal attention network (DAN), central executive network (CEN), and VN by functional labels, and in frontal, temporal, parietal, and occipital lobes by anatomical label. The *k*_max_-core voxel was classified using larger functional labels (7 ICs) (e) and anatomical labels (8 lobes) (f). In this individual, VN or parietal/occipital lobe voxels belonged in the largest number to the *k*_max_-core.

When *k*-cores derived from *k*-core percolation were embedded on a hyperbolic disc, as the *k* increased, voxels farther from the center of the disc were removed earlier, and then voxels near to the disc center tended to remain in the *k*-cores and final survivors composed *k*_max_-core (Fig 4a-b, Supplementary Figure 5, 6).

Due to the varying number of edges at the beginning, the number and location of *k*_max_-core voxels varied highly from individual to individual (mean number of voxels at *k*_max_-core: 826 ± 503, range: 251-1854). We counted how many *k*_max_-core voxels belonged to specific resting-state ICs/lobes subnetworks (Fig. 5). According to the ICs, VN1 (range: 0-244)/ VN2 (range: 0-328) / VN3 (range: 0-196), Visual Attention Network (VAN) (range: 0-258) showed the larger mean number of voxels comprising *k*_max_-core (Fig. 5a, Supplementary Table 5). Once combining ICs into seven networks, VN (sum of VN1, 2, 3, 4 and VAN) (range: 0-790) had the largest number of *k*_max_-core voxels, and DAN (range: 0-191) had the smallest (Fig. 5b, Supplementary Table 6). According to the lobes, left (range: 22-327) and right (range: 3-314) parietal and left (range: 0-236) and right (range: 0-243) occipital lobes had the larger number of *k*_max_-core voxels (Fig. 5c, Supplementary Table 7). Using eight-category anatomical label, parietal (range: 25-631) and occipital (range: 0-455) had the largest *k*_max_-core voxels as well (Fig. 5d, Supplementary Table 8).

**Figure 5.**
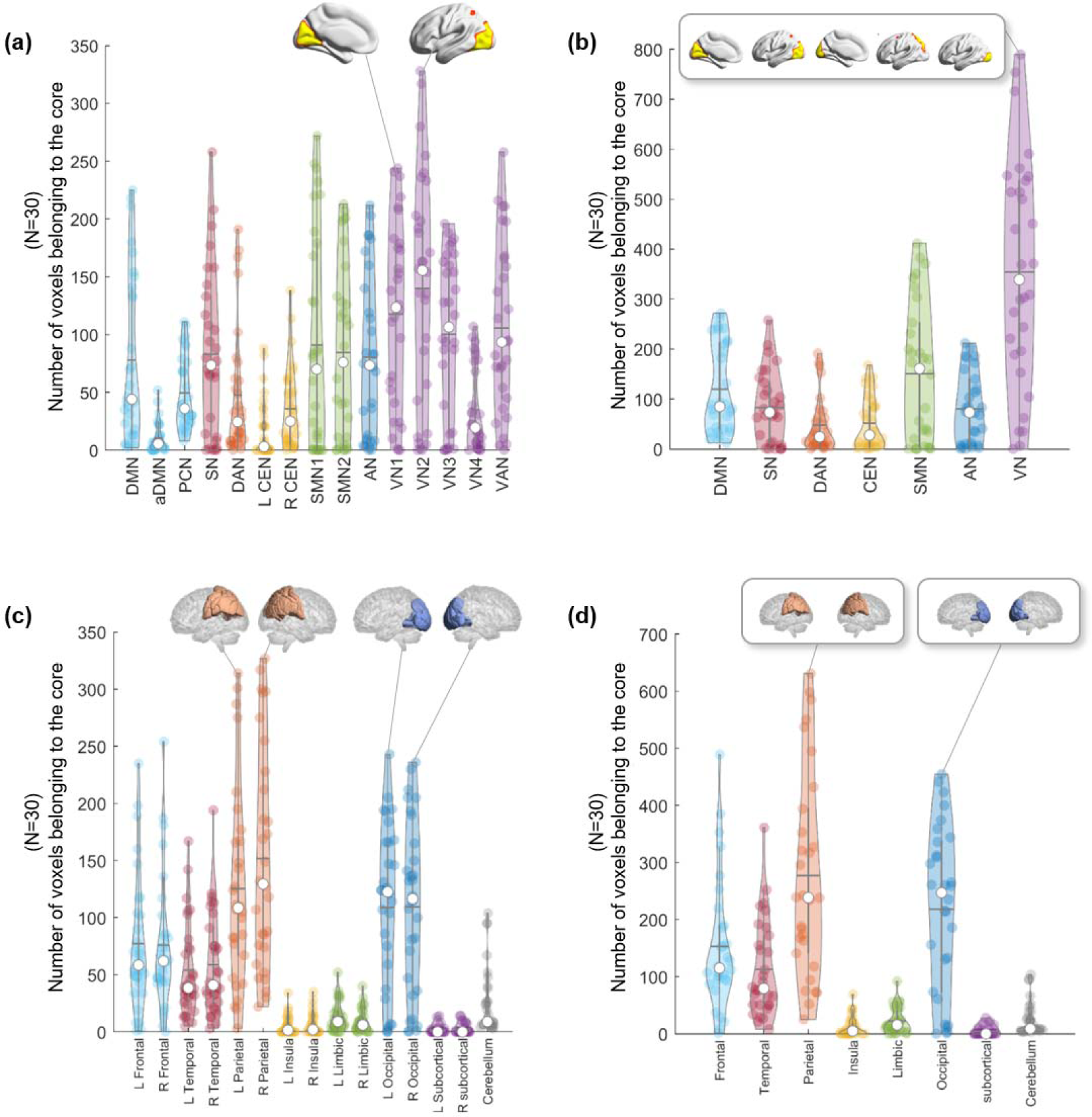
The plots show which subnetworks, in 30 individuals, the *k*_max_-core voxels belong to. *k*-core percolation yielded *k*_max_-core voxels for each individual and which independent components (ICs) or lobes those *k*_max_-core voxels belonged. ICs were represented as fifteen (a) or as the seven categorized (b). Five visual networks (VN1/2/3/4, VAN) into one visual network (VN), etc. Since some of the voxels belong to multiple ICs, categorized VN had a little bit fewer voxels than the sum of the number of voxels of constituents (V1/2/3/4, VAN). (a) According to the fifteen functional labels, visual subnetworks, VN1,2,3, and visual attention network (VAN) were leading in the number of voxels among *k*_max_-core voxels, and sensorimotor networks (SMN 1,2) and salience network (SN) followed. (b) Once categorized, the propensity of VN among the seven was outstanding. (c) Anatomical labels for both lobes and cerebellum showed prominence of occipital and parietal lobes among the fifteen, and (d) once categorized to eight, parietal and occipital lobes were sustained.

### Voxels-subnetworks composition of k_max_-core and their subnetwork distribution pattern among individuals

Voxels remaining at the last step of k-core percolation were annotated to the ICs or the lobes and their initial degrees were rendered as histogram, which revealed degree distribution of the *k*_max_-core voxels ranging from 187 to 3847 or from 21% to 52% (the percent of the degree of each *k*_max_-core voxel per the degree of the voxel with the highest degree). Interestingly, the distribution of the degrees of the *k*_max_-core voxels varied between individuals who showed a spectrum in the distribution of the dominance (or non-dominance meaning even participation of voxels to *k*_max_-core ) of the ICs to which the *k*_max_-core voxels belonged (Supplementary Figures 7-11).

More in detail, VN included the largest number of *k*_max_-core voxels in 73% of subjects (22 among 30) than any other resting-state IC subnetworks, and more than half of the *k*_max_-core voxels belonged to VN in 40% of subjects (12 among 30). The degrees of *k*_max_-core voxels belonging to VN ranged from high to low values just like the degrees of the voxels belonging to the other ICs such as DAN, DMN, SN (Supplementary Figure 10, 11). Over the individual differences, we questioned whether there was any group-level *k*_max_-core and found that 34 *k*_max_-core voxels were shared in common by more than 60% of subjects (18 among 30), and VN occupied the largest number of these common *k*_max_-core voxels (Fig. 6a, b). VN could be said to be the most dominant IC subnetwork (V1: 21, V2: 14, V3: 16, and VAN: 6 voxels).

**Figure 6.**
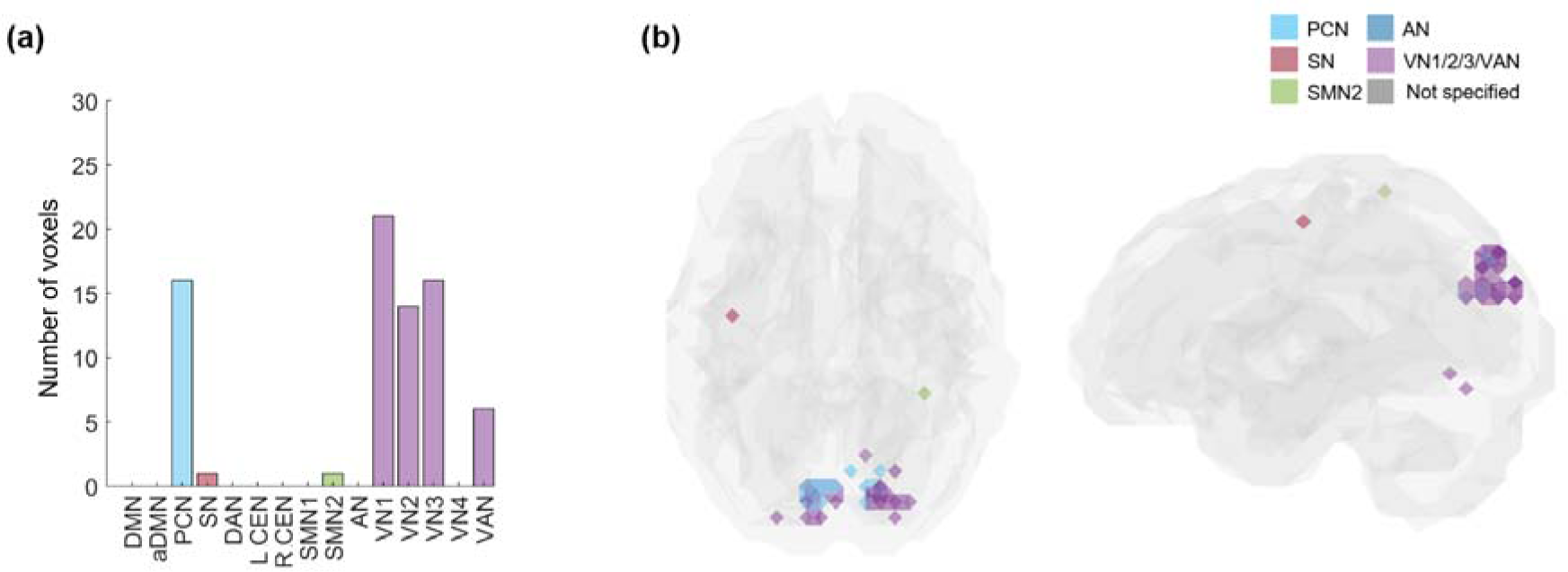
The common core voxels shared by 60% of individuals. *k*-core percolation disclosed to which IC subnetworks prevalently among individuals, the *k*_max_-core voxels belonged. (a) A bar plot is showing the affiliation of *k*_max_-core voxels. Shared voxels of precuneus network (PCN), visual network1 (VN1), VN2, V3, and visual attention network (VAN) were easily found and a rare voxel in salience (SN) and sensorimotor network2 (SMN2). (b) The shared voxels on the template brain were visualized. Voxels of *k*_max_-core are found in VN and PCN.

Next to VNs, PCN included the largest number of *k*_max_-core voxels (commonly shared in 60% of subjects: 16 voxels) (Fig. 6a, b). When the anatomical label was applied, 34 *k*_max_-core common voxels in 60% of subjects (n=18) mainly belonged to occipital and parietal regions. In more detail, 17 voxels in bilateral lateral superior occipital regions, 13 voxels in the bilateral ventromedial occipital regions, three voxels in the parietal regions (superior parietal lobule, precuneus, postcentral gyrus), one in frontal region participated in the *k*_max_-core. We suggest that posterior area voxels including VN/PCN or occipitoparietal regions are the candidates for common core in more than half of the normal subjects to preside over their hierarchically lower voxels in the same ICs and the voxels belonging to the other ICs or lobes.

We grossly found the three patterns of *k*_max_-core voxels-IC composition among individuals using functional label: DMN-dominant, VN-dominant, and uneven but distributed (Fig. 7, Supplementary Figure 11). When a dominant IC (DMN or VN) occupies more than 40% of *k*_max_-core voxels, the individuals were deemed to be DMN-dominant or VN-dominant (Figure. 7, Supplementary Figure 11a, b, 12a, b). An individual without any dominant IC and showed the distributed (*k*_max_-core) voxels-IC compositions were named as having a distributed pattern (Figure. 7c, Supplementary Figure 11c, 12c). There were five individuals with DMN-dominant pattern, 18 individuals with VN-dominant pattern, and seven individuals who had a distributed pattern (Supplementary Figure 11). The *k*_max_-core voxels’ sizes of the individuals with the distributed pattern were significantly greater than those of the individuals with VN- or DMN-dominant patterns (*p* < 0.05). Stacked histograms of the degree distribution of *k*_max_- core voxels are presented in Fig. 8 and it is to be noted that all the 30 individuals, regardless of degree maximum of each individual, could be annotated with one of the three types of *k*_max_-core voxels-ICs composition. An exemplary case showed the changes of k-cores in terms of IC-voxel rendering on brain template along k-core percolation (Supplementary Figure 13).

**Figure 7.**
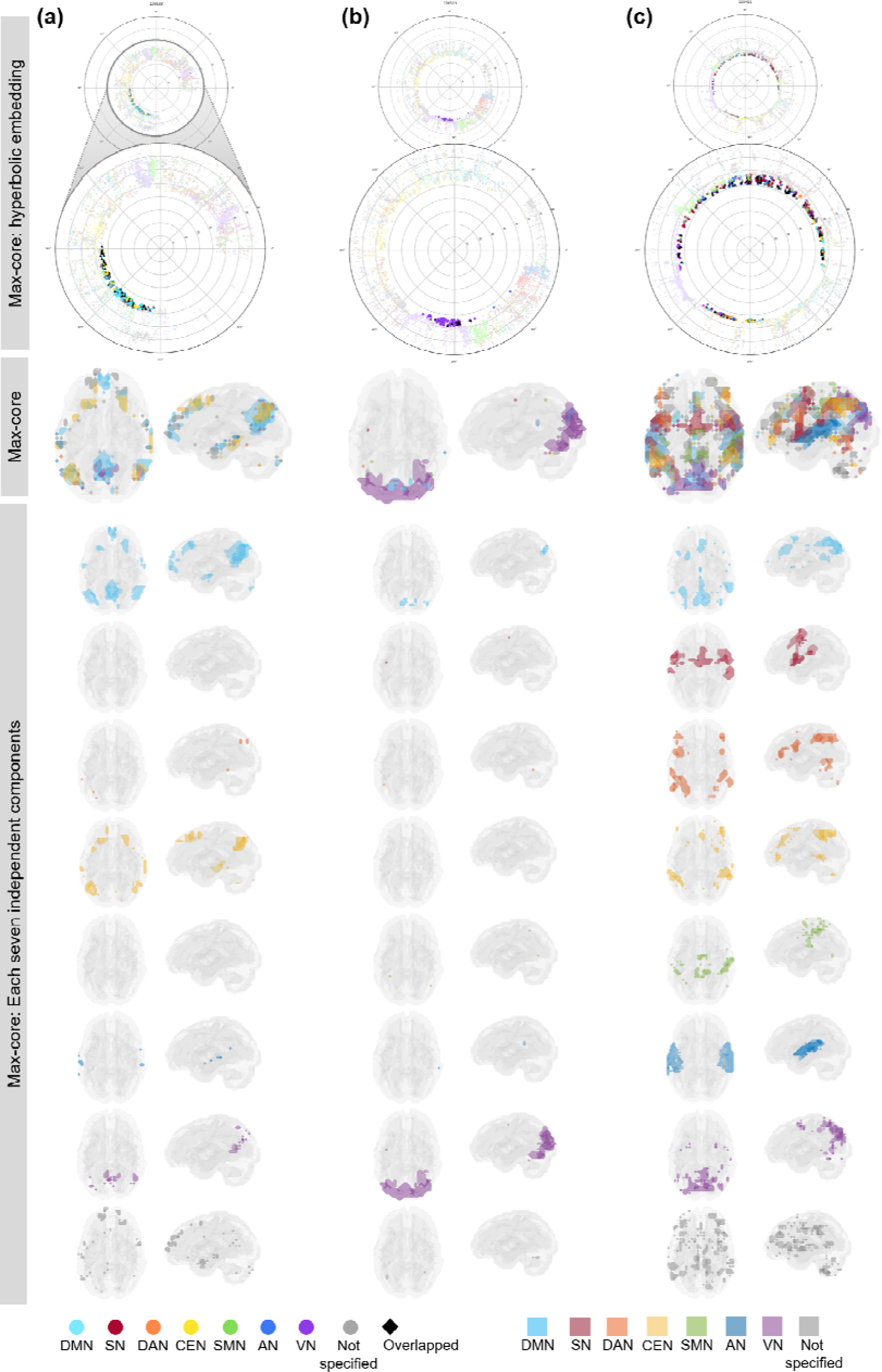
Three types of *k*_max_-core voxels-independent components (ICs) composition at the end of *k*-core percolation. There were three types, named based on which ICs the *k*_max_- core voxels belong to; VN-dominant, DMN-dominant and the distributed. Categorized functional labels, consisting of default mode network (DMN), salience network (SN), dorsal attention network (DAN), central executive network (CEN), auditory network (AN), and visual network (VN) were used to classify *k*_max_-core voxels according to their belonging to these categorized labels. In the top row, each individual’s *k* max core was embedded on the hyperbolic disc. The *k*_max_-core was enlarged and shown in detail. In the middle, *k*_max_-core were visualized on the 3-dimensional brain. At the bottom, the *k*_max_-core voxels belonging to seven networks were shown separately. (a) Individuals with more than 40% of their *k*_max_-core voxels being in DMN were classified DMN-dominant. *k*_max_-core voxels of one example (129533) of DMN-dominant type shows blue regions indicating *k*_max_-core voxels that belong to DMN. More than 60% of *k*_max_-core voxels were in DMN regions ranging over precuneus, lateral parietal cortex, and medial prefrontal regions. (b) Individuals with more than 40% of *k*_max_-core voxels being in VN were classified VN-dominant. A VN-dominant individual (126325) shows more than 80% of *k*_max_-core voxels belonged to VN ranging over medial and lateral occipital and parietal regions. (c) Individuals were classified as having the distributed pattern when there are no dominant IC subnetworks found. In an example case (110411), every subnetwork voxel contributed to less than 20% of *k*_max_-core voxels. We counted all duplicates when the *k*_max_-core voxels belonged to multiple IC subnetworks.

**Figure 8.**
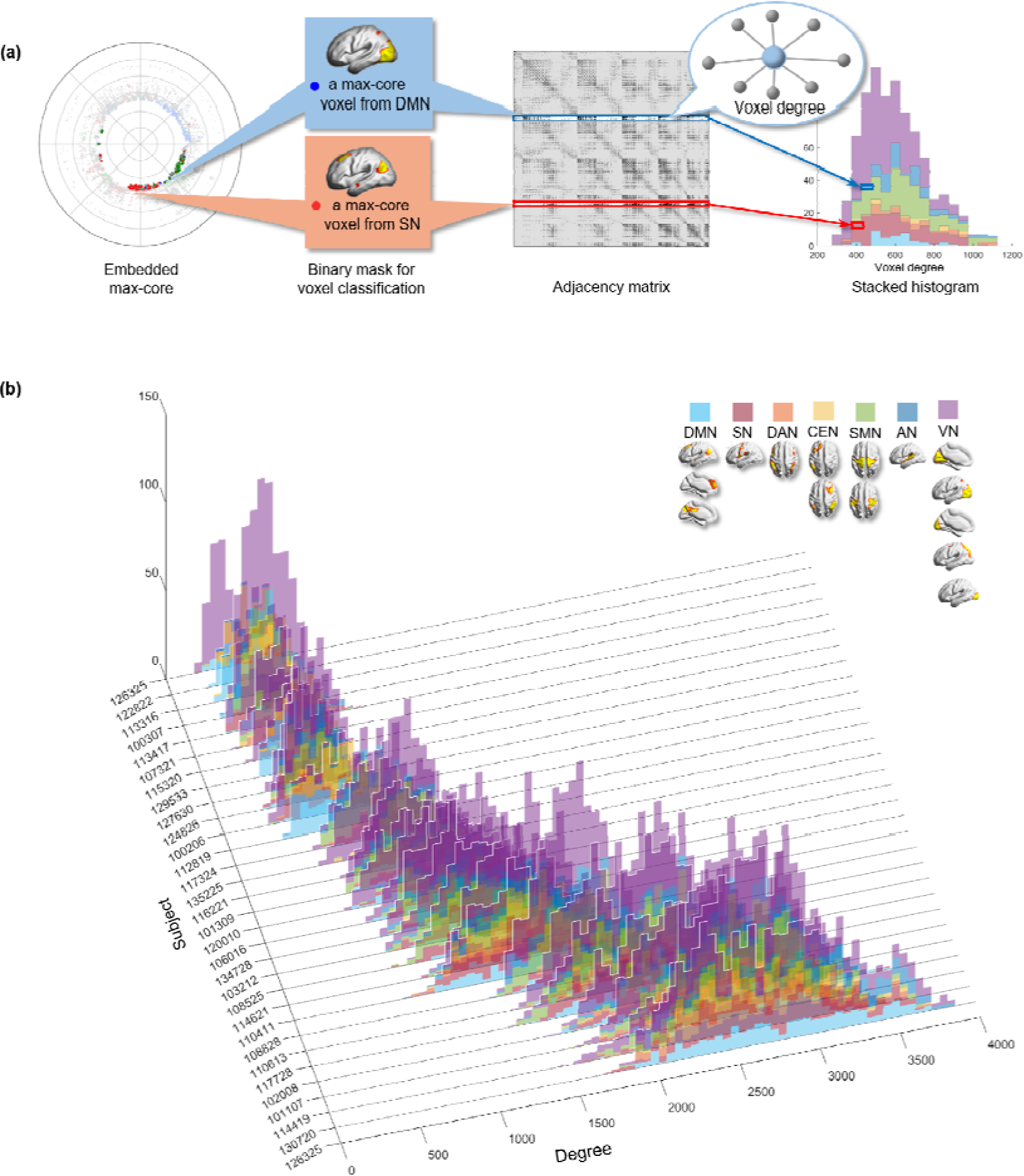
The stacked histogram of the degree distribution of *k*_max_-core voxels calculated from adjacency matrix used as input. (a) After finding *k*_max_-core of a subject with *k*-core percolation, we read the degree of each *k*_max_-core voxel on the adjacency matrix. We classified *k*_max_-core voxels into seven categorical independent components (ICs) and produced a stacked histogram showing each *k*_max_-core voxel’s affiliation and also the voxel degree simultaneously. (b) We depicted *k*_max_-core of 30 subjects. The voxel degrees of *k*_max_- core voxels and their affiliation were in different colors. *k*_max_-core voxels located on the right side of the histogram were deemed to have a greater degree initially in the adjacency matrix, indicating that they have connections with non-*k*_max_-core voxels as well as within *k*_max_-core voxels. In contrast, *k*_max_-core voxels on the left side of the histogram denote a relatively smaller voxel degree, implying it has fewer connections with non-*k*_max_-core voxels and thus almost confined to get connections with *k*_max_-core voxels. The histograms of 30 subjects were sorted in ascending order with the mean degree of the *k*_max_-core voxels in 3-dimensional space.

## Discussion

In this study, we applied hyperbolic embedding and *k*-core percolation to investigate the latent topology of the human functional brain network. The hyperbolic embedding provides insightful visualization for the tree-like nature of the functional brain network, and k-core percolation discovered the core structure of individuals. The coordinates of voxels by hyperbolic embedding enable us to measure functional within-themselves proximity of specific subnetworks of the brain. VNs and the parietal lobe showed functional proximity at rest frequently in individuals. The *k*-core percolation disclosed which subnetworks contributed to form core voxels at maximum and that there were individual variations in dominant (visual or DMN) or distributed patterns on the *k*_max_-core voxels-ICs composition analysis.

The *k*-core percolation has recently been interpreted as one part of explosive percolation [41], which was once considered to be discontinuous but later to be continuous [42] and finally proposed to be hybrid [43] in its configuration dynamics according to the percolation process. In explosive percolation in complex networks, dynamic changes of network configuration were observed/simulated in a forward way, meaning how the global connectivity was formed with the addition of new edges to the networks. The formation of the largest component could have been discontinuous, continuous or hybrid [41–43]. On the contrary, optimal percolation was also introduced to find the vulnerable nodes for targeted attack and trial of dismantling [11, 12] eventually to have defined influencer nodes [10]. This idea has come more popularly to be used in understanding epidemic spread, power grid failure [14, 15] and social message propagation [13, 25], either viral or fake, and introduced the algorithms of collective influence or belief propagation [13]. Optimal percolation tried to reveal a minimal set of nodes and thus kept itself different from *k*-core percolation decomposition saying that *k*-core percolation finds a group of nodes not pinpointing the nodes with the highest collective influencer score (for a message, electrical power, epidemic spreading capability). In optimal percolation or collective influence algorithm, of course, they proceed backward from the largest component to fragments. We followed the idea of *k*-core percolation, the largest component with the initial input and the adjacent matrix was re-configured while removing *k*-shell in this investigation.

Either forward or backward, regardless of the name and the use of specific percolation in the application context, for investigation of complex brain networks, we need to identify the voxels of highest interest and their grouping to a certain category, i.e., functional or anatomical label. When we have a sufficient number of voxels like more than 5,000 in this study, we do not need to find each voxel for its contribution maintaining resting-state brain function but want to find the groups of voxels of interest which remain after parsimonious filtration, in other words, percolation. *k*-core percolation functioned as the tool to find the survivors of this endeavor. Adoption of *k*-core percolation algorithm from the literature [27-29,44] easily yielded the voxels of *k*_max_-core. After reaching the *k*_max_, if one step further, the largest component was disintegrated into many smaller pieces which prohibited further the work of k-shell peeling in all the 180 individuals. Expectedly, the degree distribution of voxels belonging to the ICs of these *k*_max_-cores ranged from the highest to the mid-level (Supplementary Figure 9, 10).

In resting state, fMRI renders the information of the fluctuation of BOLD signals per each voxel and evidently, the individuals are conscious, though sometimes their minds are drifting from introspection/ imagination to paying attention to the milieu, full of MRI radiofrequency-derived noise or the scenes within the gantry, etc. Individual difference of this mind-setting is well expected, and also the temporal fluctuations will add up to make the stationary setting. In this study, we assumed perfect stationarity, meaning we just calculated a single-digit correlation between every voxel *i* and voxel *j*, and this led us to define 25 million or more possible edges. After scrutinizing the correlation, which ranged from -0.7 to 0.8 in most cases, by using scale-free criteria, we cut off the edges not reaching the threshold, which was 0.4.

This yielded around 300,000 to 3,900,000 edges with more than 80% of voxels remaining among 5,937 re-sized voxels. Only the positive inter-voxel correlation was brought into the analysis in this study and the negative correlation (anticorrelation) and its contribution to make multi-layered duplex interdependent brain network remains to be studied. Such is also the case with the temporal variation of *k*_max_-core voxels and their ICs-voxels composition.

Using the adjacency matrix and its consequent network, which defined the stationary voxels correlations at rest on rsfMRI, we proceeded to visualize the configuration on 2-dimension with the hyperbolic disc. All the voxels were designated to belong to one (rarely more than one) ICs and the behavior of dynamic change of surviving voxels at each step of *k*-core percolation was presented using flags-plots (Fig. 3c, Fig. 4c,d) and also on the embedded hyperbolic discs (Fig. 4a, Supplementary Fig. 5a,b). Flags-plot of voxels belonging to seven large representative ICs were displayed in Supplementary Figure 7. It was interesting that the size of voxels varied among ICs before *k*-core percolation, however, the *k*-core percolation let the certain IC-voxels such as SMN, auditory network (AN), or central executive network (CEN) vanish. Another point of merit is that voxels belonging to SN disappear completely in a few individuals but remain in the others. Regarding DAN, CEN, AN, voxels seemed to vanish, but a small number of voxels belonging to these ICs remained definitely and join the group of voxels of *k*_max_-core. This interesting phenomenon on *k*-core percolation is just reported here and will be the subject of further study to understand the conscious resting-state of mind in normal individuals and its correlates on rsfMRI.

Conscious individuals, evidently at awake resting state on examinations, and their electrophysiological or perfusion correlates on electroencephalography (EEG), magnetoencephalography (MEG) or rsfMRI were studied by various investigators to yield a representative theory of consciousness such as global neuronal workplace theory and integrated information theory (IIT) [45–50]. Global neuronal workplace theory [46, 49] advocated the distributed subnetworks interconnected onto each other at conscious states.

Thus the isolated subnetworks are not the correlates for consciousness, and instead, once connected in a network of subnetworks, the consciousness is achieved, emphasizing input-output processing. On the contrary, integrated information theory [47, 48] measures cause-effect power with maximally irreducible integrated information on some area, suggesting most likely the posterior area of the brain. Irrespective of which theory suits the data better, the subnetworks participating in the maintenance of consciousness should be discovered on each modality (EEG, MEG, or rsfMRI). If it is true that the posterior area is the one to contribute to the conscious resting-state, as indicated by IIT, we need to investigate whether the VN we found in this study would be in charge. Perturbation study and/or calculation of integrated information, ϕ, of IIT will also be necessary [50]. The method we introduced in this study will be a good platform for visualization of inter-voxel correlations and for elucidation of the changes of IC-voxels composition upon *k*-core percolation using a flags-plot for these studies.

Discovery that the *k*_max_-core voxels of the parietooccipital area are dominant in more than half of the studied individuals and that VNs participated obviously in the *k*_max_-core in the remaining distributed or DMN dominant individuals would mean that posterior or parietooccipital areas are one of the important correlates of resting-state consciousness. This indicates that everyone had VN in their *k*_max_-core voxels (Supplementary Figures 7 and 10). Interestingly, in the individuals with *k*_max_-core voxels uneven-but-distributed belonging to all the ICs, the contribution as core structure by all the other ICs seemed equivalent. The meaning of this phenomenon might be understood by looking into the temporal fluctuation of *k*_max_-core voxels-ICs investigation in the next study.

Referring to the high angular coherence on the hyperbolic disc embedding and also the dominance for *k*_max_-core on *k*-core percolation, the VNs had the strongest connectivity within IC and their dominance in *k*_max_-core voxels were significant in half of the individuals at rest. The VN has been reported as one of the major hub regions in previous studies [51]. The VN is a unimodal area that conducts highly specialized functions; for example, the primary visual cortex, well delineated by the cytoarchitectonic features is an important correlate of corresponding function vision. In contrast, the domain-general frontal area, involved in various cognitive tasks [52], showed low functional proximity within itself. The VN also has a higher neuronal density than the others. Considering that the resting functional connectomes are not engaged at any activation tasks, the unimodal highly differentiated subnetwork might be the one with strength in angular coherence and contribution to the coreness on *k*-core percolation. In the same vein, a sensorimotor network also showed high angular coherence. But it was not found to be dominant in any individual but just one of the components weakly or absent contributor to the *k*_max_-core (Supplementary Figure 7)

The voxels belonging to the precuneus network remained in the *k*_max_-core in most of the individuals. Precuneus is an associative region, especially involved in self-related information processing [53, 54]. Precuneus is also a crucial component of DMN, which was also found as dominant IC subnetwork in a fraction of the individuals in this study, reminding that DMN is the well-known subnetwork active at resting state. Though the precuneus is not inherently highly differentiated and specialized like a visual system, at resting-state, the voxels of precuneus came to join the *k*_max_-core.

The core of the individuals showed diversity in size and IC-voxels composition. This IC- voxels composition pattern was arbitrarily classified among the individuals: DMN-dominant, VN-dominant, and uneven but distributed types. Individuals having DMN or VN-dominance revealed that their *k*_max_-core voxels consisted of 1) mainly DMN and several minor ICs or 2) VN and CEN and few minors at resting-state, respectively. This might imply that dominant IC subnetworks sustain as characteristics of the individuals, and/or that in every individual, there is a fluctuation of mental states, which are drifting between the distributed pattern and, in some instances, entering the DMN major or the VN major states which would also be observed by rsfMRI. Former or latter, whichever interpretation might be the fact/truth, further study with sliding window segmentation and phase-shift observation of rsfMRI is warranted.

Individuals with the distributed pattern tended to have a larger size of voxels in their *k*_max_- core, and more than half of all had the voxels belonging to VNs or DMN in their *k*_max_-core. It is really interesting to know whether DMN or VN-dominant patterns might be a drifting accentuation of these two subnetworks while evenly distributed IC-voxels are the background default state at rest in humans. Then, DMN and VN-dominant patterns are the two extremes of the spectrum, and the distributed pattern resides between them. Regarding hubs, there was a controversy that in some reports [55, 56], hub regions were composed of voxels belonging to various subnetworks and not confined to a specific system in one report. In the other reports [57], there was the predominance of the visual system and precuneus among hub nodes. We suggest that this *k_max_* core IC-voxels composition be used as a fingerprint to identify and describe an individual’s physiological or pathological characteristics of their resting IC compositions of the cores.

Finally, finding *k_max_* core voxels using an established visualization method on hyperbolically embedded discs accompanying the *k*-core percolation raised the possibility of studying stationary and dynamic functional connectivity of voxels and their hierarchy upon filtration/ percolation. The *k*-core percolation disintegrated the initial largest component gradually but sometimes abruptly and thus this descent pattern seemed to represent core and sub-core configuration of voxels hidden under just the simple-looking scale-free functional brain network. Hidden relation between areas/ regions/ voxels on rsfMRI was recently investigated either coactivation pattern (CAP) [58] or hidden Markov model (HMM) [59, 60], both of which followed the success of elucidation of dynamic changes of various CAPs on the analysis of MEG data [61] or of discovering variable HMM states and finding the transition between HMM states at rest using MEG [62], respectively. On the temporal scale and/or cross-modal investigations, both CAP and HMM methods were used to understand the twitches and other trivial movement/activities of humans during imaging [63] and eventually consciousness. Our method of *k*_max_-core detection and the annotation of the *k*_max_-core voxels to ICs upon filtration will lead us to define the hierarchical structure of core-periphery coherent gathering of voxels. The 2-dimensional display on hyperbolic discs let us visualize how *k*_max_-core was formed by the simple rule of *k*-shell peeling or decomposition. This is the new platform to understand finally the awake, twitching intermittently, mentally drifting with various attention to milieu or his/her mind in the MRI gantry in a conscious resting-state of human individuals. Temporal variation [64] with rsfMRI and cross-modal investigation with MEG or EEG [65] will be the next step of investigation using the current method of hyperbolic disc embedding and *k*-core percolation.

## Method

### Dataset

We included 180 participants from the Human connectome project (HCP) S1200 release, which is available with open access (www.humanconnectomeproject.org). The acquisition parameters and preprocessing steps were described in [66]. Participants were free of neurological diseases and psychiatric disorders (31 participants with age range 22-25 years, 84 participants with age range 26-30 years, and 64 participants with age range 31-36 years; male: 76, female: 104). We included the whole 180 subjects in angular coherence analysis, and 30 subjects in *k*-core percolation analysis (ten participants with age range 22-25 years, ten participants with age range 26-30 years, and ten participants with age range 31-36 years; male: 15, female: 15).

### Preprocessing of the rsfMRI data

The rsfMRI was obtained with 3T scanner with following parameter: TR = 720 ms; TE = 33.1 ms; flip angle = 52°; FOV = 208 × 180 mm, 2 mm isotropic voxels. The minimally preprocessed data from HCP was further preprocessed using Statistical Parametric Mapping (SPM, www.fil.ion.ucl.ac.uk/spm/) and FMRIB Software Library (FSL, fsl.fmrib.ox.ac.uk/fsl/) [66]. We corrected gradient and motion-induced distortion, and field map-based non-linear transform was also used to correct distortion. After images were coregistered and normalized into standard space, intensity normalization was done. These minimal preprocessing results in 2 × 2 × 2 mm sized voxel images [66]. We additionally conducted smoothing using 6 mm full-width at half maximum (FWHM) of the Gaussian kernel, and the band-pass filtering (0.01 Hz-0.1 Hz). Finally, we down-sampled the data to reduce the computational load of voxel-based whole-brain network analysis (dimension: 31 × 37 × 31, voxel size: 6 × 6 × 6 mm ), and 5,937 voxels that consisted of the gray matter were used.

### Functional/anatomical label of voxels

We performed Independent Component Analysis (ICA) using MELODIC (Multivariate Exploratory Linear Optimized Decomposition into Independent Components) to extract resting-state networks [36]. Fifteen independent components were classified after manual inspection of spatial maps: aDMN, DMN, PCN (equivalent to posterior DMN), SN, DAN, left CEN (L CEN), right CEN (R CEN), SMN1, SMN2, AN, VN1, VN2, VN3, VN4, VAN. The spatial maps were manually inspected and classified based on previous studies [67, 68]. To present our results in a more comprehensible summary, we also combined 15 resting-state networks into seven categories: DMN, SN, DAN, CEN, SMN, AN, VN. Both schemes were used for functional label. Brainnetome atlas was used to generate 15 left/right anatomical lobes, and also seven lobes consisted of bilateral brain regions for anatomical label [40].

### Assessment of functional connectivity and voxels-composition of subnetworks

For the scale of the 5,937 cubic isotropic voxels, we measured blood oxygen level dependent (BOLD) signals from each voxel of fMRI data and characterized the spontaneous fluctuations over the time-series, σ^2^(X), where X is the time-series of BOLD signal. For the BOLD-fMRI time-series X = (X_1_, …, X_N_) of a given voxel, the variance was computed by the sample variance σ^2^(*X*).. given as the following formula:

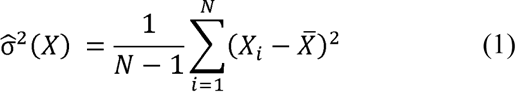

where *X̄* denoted the sample mean of 1.. Between a pair of voxels with BOLD-fMRI time-series X = (X_1_, …, X_N_) and Y = (Y_1_, …, Y_N_), the functional connectivity was estimated by the sample Pearson correlation coefficient ρ̂:

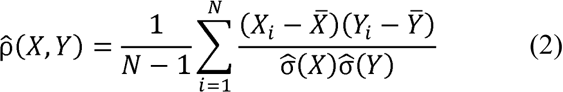

From sample Pearson correlation coefficient ρ̂ of each pair of voxels, we obtained a square matrix of Pearson correlation coefficient for each of the subjects. For determination of the most appropriate threshold values for composing a binary network, we applied tentative threshold values for ρ̂, followed by comparison of the degree distribution and the relative size of the largest connected component. We determined the threshold value, considering both the scale-freeness of degree distribution and the maximal inclusion of voxels as possible in the largest component. For scale-freeness, we considered degree distribution with a straight- formed line on a log-log scale plot as appropriate. For the size of the network, we considered the threshold low enough for the single largest component of a graph (which was embeddable on the disc) to contain 80% of the voxels in the brain. Consequently, we constructed an unweighted, undirected graph for each subject by applying the threshold to the correlation coefficient matrix.

Hyperbolic disc embedding of networks into 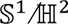 model

The resulting binary graph from each subject was mapped onto a hyperbolic disc, using 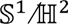 geometric network model [31, 34]. Connectomes were assumed to exist in underlying geometric space, linked with the observed topologic properties through a law of connection probability that defined the likelihood if the two regions of the brain were linked. In the model, the connection probability between two voxels, and I was determined by the hyperbolic distance between two voxels:

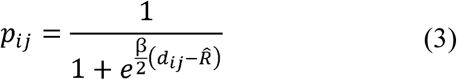

where β was the clustering coefficient of network and *R̂* was the outermost radial coordinate among the embedded voxels. The hyperbolic distance, *d*_*ij*_, was usually determined by hyperbolic laws of cosines,

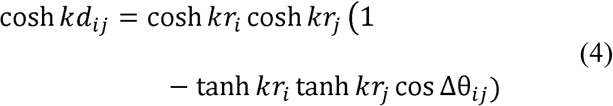

where *k* was the curvature of plane, *r*_*j*_ was the radial coordinate of the ith point and Δθ was the angular separation between two points i and j, while,

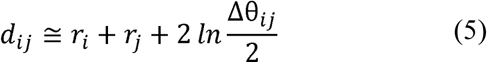

was used as a good approximation^1^. To determine the geometric object (i.e., a hyperbolic disc) that was most likely to generate the original binary graph, we implemented a software named *Mercator*^2^, introduced by García-Pérez et al. [31], which applied maximum likelihood techniques and machine learning approach for embedding of the network. By this means, by computing the most appropriate set of polar coordinates for *N* voxels (r_1_, θ_1_), (r_2_, θ_2_), (r_*N*_, θ_*N*_) and the clustering coefficient β, we embedded the binary graph onto a hyperbolic disc for each subject.

### Angular coherence to assess the degree of gathering of subnetwork voxels on the hyperbolic discs

For the assessment of narrow or wide-spread aggregation of voxels in the hyperbolic disc space, we used a metric called angular coherence, which was previously devised to investigate how points were angularly similar on the hyperbolic disc [32]. The angular coherence ξE l0,1 of a set of point 1., was determined as follows:

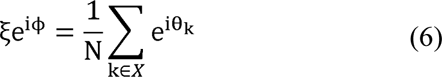

where θ is coordinate of the point and N was the number of points (voxels in this study) in *X*. As intuitive from Eq. (6) the higher ξ was, the more points in the set were locally concentrated to form an angularly gathered structure. In this study, we used binarized (Z > 6)

IC maps for functional label and Brainnetome atlas for anatomical label. The degree of gathering of subnetworks on hyperbolic discs for resting-state voxels networks was estimated by calculating the angular coherences of the voxels included in the ICs or lobes.

### k-core percolation and composition of k-core and k_max_-core structures

We performed *k*-core percolation to investigate individual-specific core subnetworks of brain connectivity. The *k-core* of a network meant the maximal subgraph of the network in which all vertices had degree at least . The -core was identified by removing all voxels with degree less than, recalculating the degrees of all the remaining voxels, until no voxel remains with the degree less than . We used a pruning algorithm suggested by Azimi-

Tafreshi et al [29]. The voxels that belonged to the -core but not to the ck 1.-core form the -shell of the network, and they were said to have -coreness. Along *k*-core percolation, the changes of the size of subnetwork at *k*-core was well visualized in these flags-plots over increasing *k,* revealing the association between the coreness of voxels and subnetworks of ICs or lobes that the voxels belonged to. Voxels and their belonging to ICs or lobes were drawn as flags-plot to describe how the number of voxels consisting of each IC/lobe decreased over increasing coreness *k*.

### Degree distribution of k_max_-core over functional/anatomical subsystems

As the coreness *k* increased, step maximum was defined as the one where on step further, at *k*_*max*_ + 1, all voxels constituting the largest component were disintegrated to smaller pieces. This phase transition happened always abruptly and thus discontinuous which made *k*_max_-core.

We evaluated the size and degree distribution of this *k*_max_-core for ICs and lobes.

## Data availability

The functional brain dataset in the present study is available for download on the Human Connectome Project (HCP) (https://www.humanconnectome.org/) with the acceptance of data use terms.

## Acknowledgments

This work was supported by the National Research Foundation of Korea (NRF) grant funded by the Korean Government (MSIP) (no. 2017M3C7A1048079, no. 2020R1A2C2101069, no. 2020R1A2C2011532 and no. 2017R1A5A1015626).

## Ethics declarations

### Conflict of Interest

The authors declare no conflict of interest.

### Supplementary Information

Supplementary Tables 1-8

Supplementary Figures 1-13

## Footnotes

1 This approximation is used for Δθ_*ij*_ that is not too small, for 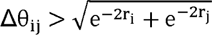

2 Available at https://github.com/networkgeometry/mercator

## Notes

### Competing Interest Statement

The authors have declared no competing interest.

### Summary of Updates

Special symbols displayed incorrectly have been fixed.

