## Supplementary Tables 1-8, Supplementary Figures 1-13 for "Characteristic core voxels in normal individuals revealed by hyperbolic disc embedding and *k*-core percolation on resting state fMRI"

**Supplementary Table 1. The average and standard deviation (SD) of angular coherence (AC) of ICs from repeated embedding.** In a subject (100206), hyperbolic disc embeddings were repeated 100 times, and ACs were calculated for IC subnetworks: default mode network (DMN), anterior DMN (aDMN), salience network (SN), dorsal attention network (DAN), left and right central executive network (L/R CEN), sensorimotor network 1/2 (SMN1/2), auditory network (AN), visual network 1/2/3/4 (V1/2/3/4), and visual attention network (VAN).

| IC | Mean | SD | CV |
| --- | --- | --- | --- |
| DMN | 0.72 | 0.06 | 0.08 |
| aDMN | 0.54 | 0.10 | 0.19 |
| PCN | 0.80 | 0.04 | 0.05 |
| SN | 0.62 | 0.01 | 0.02 |
| DAN | 0.33 | 0.12 | 0.36 |
| L CEN | 0.55 | 0.07 | 0.12 |
| R CEN | 0.57 | 0.01 | 0.02 |
| SMN1 | 0.78 | 0.05 | 0.06 |
| SMN2 | 0.62 | 0.01 | 0.01 |
| AN | 0.81 | 0.01 | 0.01 |
| VN1 | 0.84 | 0.02 | 0.02 |
| VN2 | 0.46 | 0.21 | 0.47 |
| VN3 | 0.53 | 0.08 | 0.15 |
| VN4 | 0.78 | 0.01 | 0.02 |
| VAN | 0.57 | 0.18 | 0.31 |

IC: independent component, SD:standard deviation, CV:Coefficient of variation

**Supplementary Table 2. Demographic information of subjects.** In angular coherence analysis, 180 subjects were included. Thirty subjects were included in *k*-core percolation analysis.

|  | | n=180 | n=30 |
| --- | --- | --- | --- |
| Age |  |  |  |
|  | 22-25 | 31 | 10 |
|  | 26-30 | 84 | 10 |
|  | 31-35 | 64 | 10 |
|  | 36+ | 1 | - |
| Gender |  |  |  |
|  | Male | 76 | 15 |
|  | Female | 104 | 15 |

**Supplementary Table 3. Angular coherence of each independent component (IC).** SD: standard deviation

| IC | Mean | SD | Median | Number of voxels |
| --- | --- | --- | --- | --- |
| DMN | 0.57 | 0.14 | 0.58 | 441 |
| aDMN | 0.64 | 0.18 | 0.66 | 158 |
| PCN | 0.48 | 0.15 | 0.50 | 224 |
| SN | 0.65 | 0.17 | 0.69 | 351 |
| DAN | 0.50 | 0.18 | 0.51 | 363 |
| L CEN | 0.75 | 0.16 | 0.79 | 285 |
| R CEN | 0.66 | 0.15 | 0.69 | 428 |
| SMN | 0.81 | 0.15 | 0.86 | 294 |
| SMN | 0.75 | 0.13 | 0.78 | 267 |
| AN | 0.63 | 0.16 | 0.66 | 289 |
| VN1 | 0.71 | 0.15 | 0.74 | 286 |
| VN2 | 0.72 | 0.15 | 0.76 | 443 |
| VN3 | 0.80 | 0.18 | 0.86 | 253 |
| VN4 | 0.75 | 0.22 | 0.84 | 205 |
| VAN | 0.56 | 0.16 | 0.58 | 321 |

**Supplementary Table 4. Angular coherence of each lobe.** SD: standard deviation

| Lobes | Mean | SD | Median | Number of voxels |
| --- | --- | --- | --- | --- |
| L Frontal | 0.29 | 0.13 | 0.29 | 830 |
| R Frontal | 0.30 | 0.13 | 0.30 | 820 |
| L Temporal | 0.29 | 0.12 | 0.28 | 521 |
| R Temporal | 0.25 | 0.12 | 0.23 | 495 |
| L Parietal | 0.25 | 0.12 | 0.24 | 526 |
| R Parietal | 0.26 | 0.13 | 0.24 | 549 |
| L Insula | 0.57 | 0.20 | 0.57 | 58 |
| R Insula | 0.62 | 0.19 | 0.62 | 58 |
| L Limbic | 0.40 | 0.15 | 0.39 | 115 |
| R Limbic | 0.40 | 0.17 | 0.40 | 86 |
| L Occipital | 0.71 | 0.19 | 0.76 | 320 |
| R Occipital | 0.72 | 0.19 | 0.78 | 327 |
| L Subcortical | 0.54 | 0.23 | 0.55 | 212 |
| R subcortical | 0.48 | 0.21 | 0.47 | 207 |
| Cerebellum | 0.36 | 0.18 | 0.35 | 813 |

**Supplementary Table 5. The mean numbers of *k*_max_-core voxels of 30 subjects were calculated using functional labels.** Fifteen resting-state independent component (IC) networks were used. SD: standard deviation

| IC | Mean | SD | Total voxels  belonging to IC |
| --- | --- | --- | --- |
| DMN | 78 | 73 | 441 |
| aDMN | 10 | 13 | 158 |
| PCN | 50 | 32 | 224 |
| SN | 83 | 71 | 351 |
| DAN | 48 | 56 | 363 |
| L CEN | 18 | 25 | 285 |
| R CEN | 36 | 36 | 428 |
| SMN1 | 91 | 95 | 294 |
| SMN2 | 85 | 77 | 267 |
| AN | 80 | 72 | 289 |
| VN1 | 118 | 81 | 286 |
| VN2 | 140 | 100 | 443 |
| VN3 | 100 | 70 | 253 |
| VN4 | 35 | 36 | 205 |
| VAN | 106 | 74 | 321 |

**Supplementary Table 6. The mean numbers of *k*_max_-core voxels of 30 subjects were calculated using functional labels.** Seven categories combining 15 independent components (ICs) were used. SD: standard deviation

| IC | Mean | SD | Total voxels  belonging to categorized IC |
| --- | --- | --- | --- |
| DMN | 120 | 90 | 732 |
| SN | 83 | 71 | 351 |
| DAN | 48 | 56 | 363 |
| CEN | 52 | 53 | 682 |
| SMN | 151 | 141 | 483 |
| AN | 80 | 72 | 290 |
| VN | 354 | 228 | 1104 |

**Supplementary Table 7. The mean numbers of *k*_max_-core voxels of 30 subjects for 15 anatomical labels.** SD: standard deviation

| Lobes | Mean | SD | Total voxels  belonging to lobe |
| --- | --- | --- | --- |
| R Frontal | 77 | 60 | 830 |
| L Frontal | 76 | 58 | 820 |
| R Temporal | 54 | 41 | 521 |
| L Temporal | 59 | 45 | 495 |
| R Parietal | 125 | 86 | 526 |
| L Parietal | 152 | 97 | 549 |
| R Insula | 6 | 8 | 58 |
| L Insula | 7 | 9 | 58 |
| R Limbic | 13 | 13 | 115 |
| L Limbic | 9 | 10 | 86 |
| R Occipital | 109 | 72 | 320 |
| L Occipital | 109 | 76 | 327 |
| R Subcortical | 2 | 4 | 212 |
| L Subcortical | 2 | 4 | 207 |
| Cerebellum | 26 | 30 | 813 |

**Supplementary Table 8. The mean numbers of *k*_max_-core voxels of 30 subjects for categorized anatomical labels.** SD: standard deviation

| Lobes | Mean | SD | Total voxels  belonging to each bilateral lobes |
| --- | --- | --- | --- |
| Frontal | 153 | 117 | 1650 |
| Temporal | 113 | 84 | 1016 |
| Parietal | 277 | 180 | 1075 |
| Insula | 12 | 17 | 116 |
| Limbic | 23 | 23 | 201 |
| Occipital | 218 | 146 | 647 |
| Subcortical | 4 | 7 | 419 |
| Cerebellum | 26 | 30 | 813 |

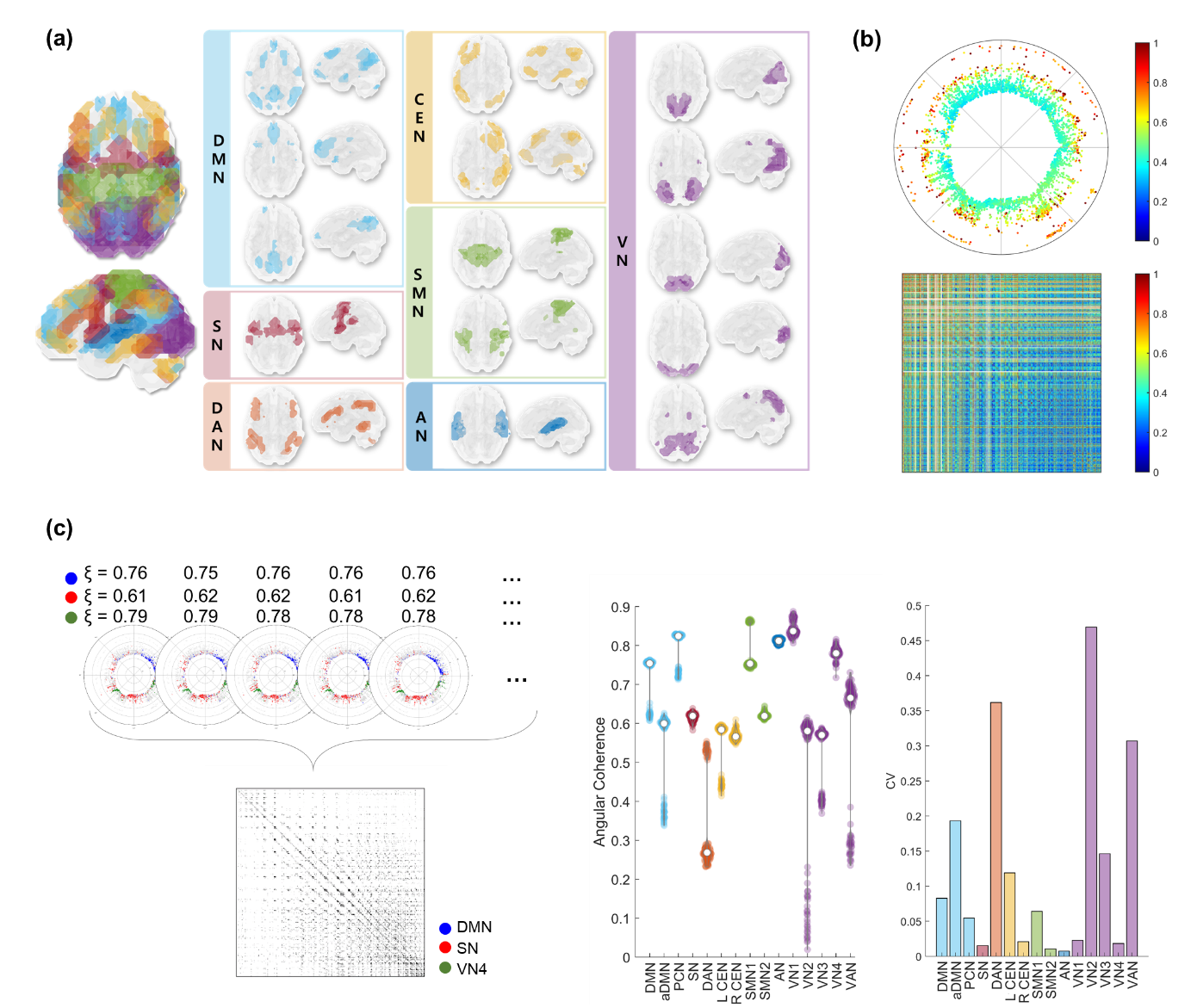

**Supplementary Figure 1. Summary of the methods; hyperbolic disc embedding and its reproducibility and *k*-core percolation with rendering upon the hyperbolically embedded discs.** (a) Fifteen independent components (ICs) derived from the independent component analysis were rendered in the 3D brain. The Default mode network (DMN), anterior DMN (aDMN), and precuneus network (PCN) were printed in pale blue, salience network (SN) in red, and dorsal attention network (DAN) in orange. The left/right central executive network (L/R CEN) were shown in yellow, sensorimotor networks 1/2 (SMN) in green, auditory network (AN) in navy blue, and visual networks (V1/2/3/4/VAN) in violet. All ICs were shown in both axial and sagittal views. (b) The reproducibility of hyperbolic embedding was shown for one example individual (subject #100206) with 100 repetitions. Using this repeatedly embedded 5,937 voxels 100 times, and the intervoxel distances in the hyperbolic disc were considered edges. After thresholding, voxels not belonging to the largest component and edges not found valid were shown in white in the matrix. The coefficient of variation (CV) of intervoxel distances were shown as a matrix (bottom). For a voxel, all the CVs of distances of its edges connecting with all the other valid edges were averaged to yield its representative CV of embedding reproducibility. On one embedded disc, CVs per voxels were depicted with colors of the jet color map. CVs per voxels ranged from 0.29 to higher values, and the voxels near the center were found to be put (embedded) reproducibly and the voxels outside to the periphery of the disc showed higher variability of the intervoxel distances with all the other voxels (their locations on the repeated embedding moves around the axis angle and thus the distance). (c) We also calculated angular coherence (AC) of each embedding using 15 ICs (left), and the distribution of AC for each IC was displayed (right). The CV of each IC was shown in a bar plot (right). Be noted that the reproducibility of AC over repeated hyperbolic disc embedding shows the reproducibility of IC-voxels configuration upon the embedded disc despite rotation/reflection symmetry and branch permutation symmetry of hyperbolic disc owing to its geometry characteristics.

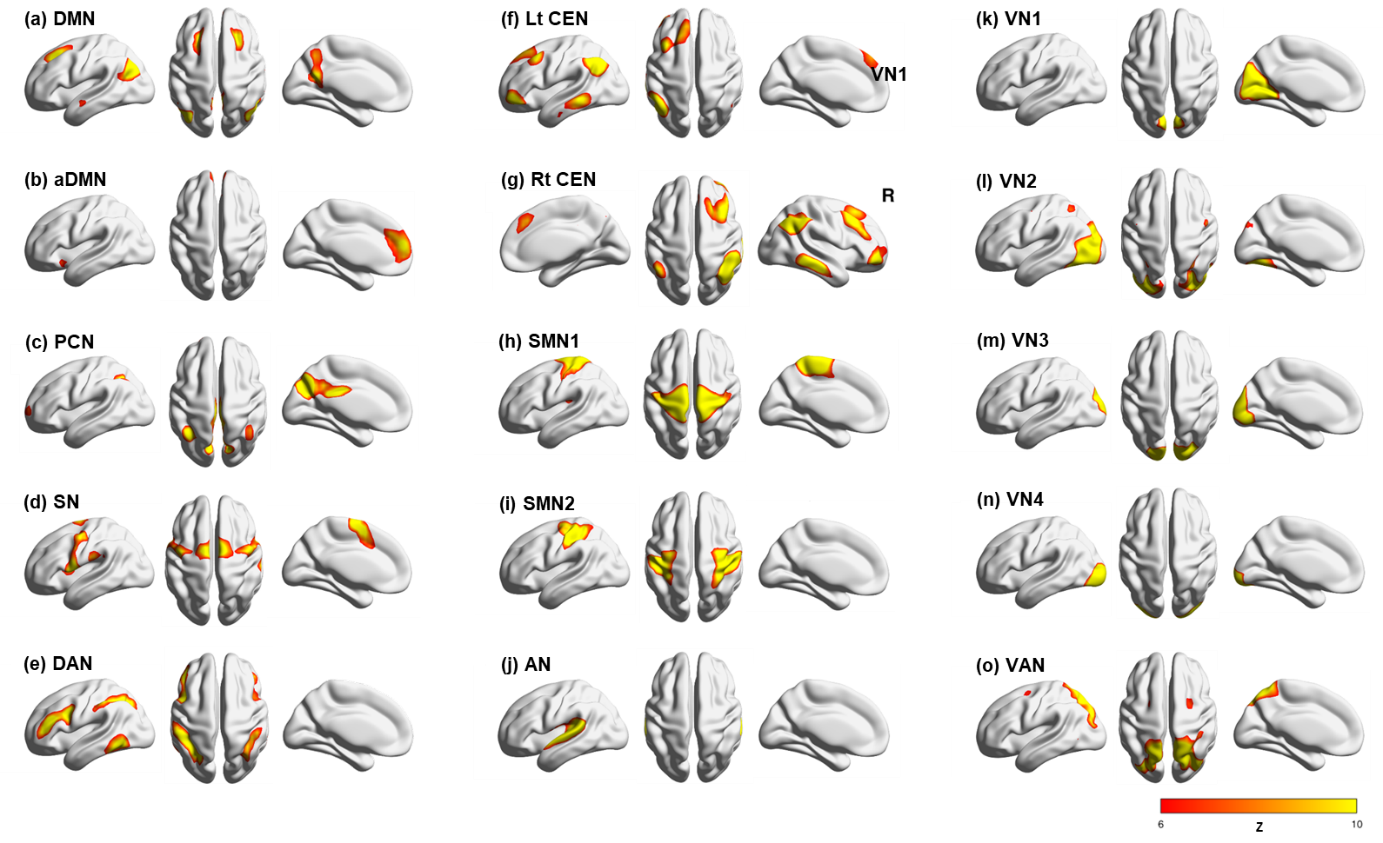

**Supplementary Figure 2. Fifteen independent component (IC) subnetworks and their functional labels.** 180 individuals’ rsfMRI data were preprocessed and resampled into 6x6x6 mm^3^ and were put into group independent component analysis. The spatial maps of ICs were illustrated on the brain surface (Z > 6). (a) Default mode network (DMN), (b) anterior DMN, (c) precuneus network (PCN), (d) salience network (SN), (e) dorsal attention network (DAN), (f) left central executive network (L CEN), (g) right CEN (R CEN), (h) sensorimotor network 1 (SMN1), (i) SMN 2, (j) auditory network (AN), (k) visual network 1 (VN1), (l) VN2, (m) VN3, (n) VN4, (o) visual attention network (VAN).

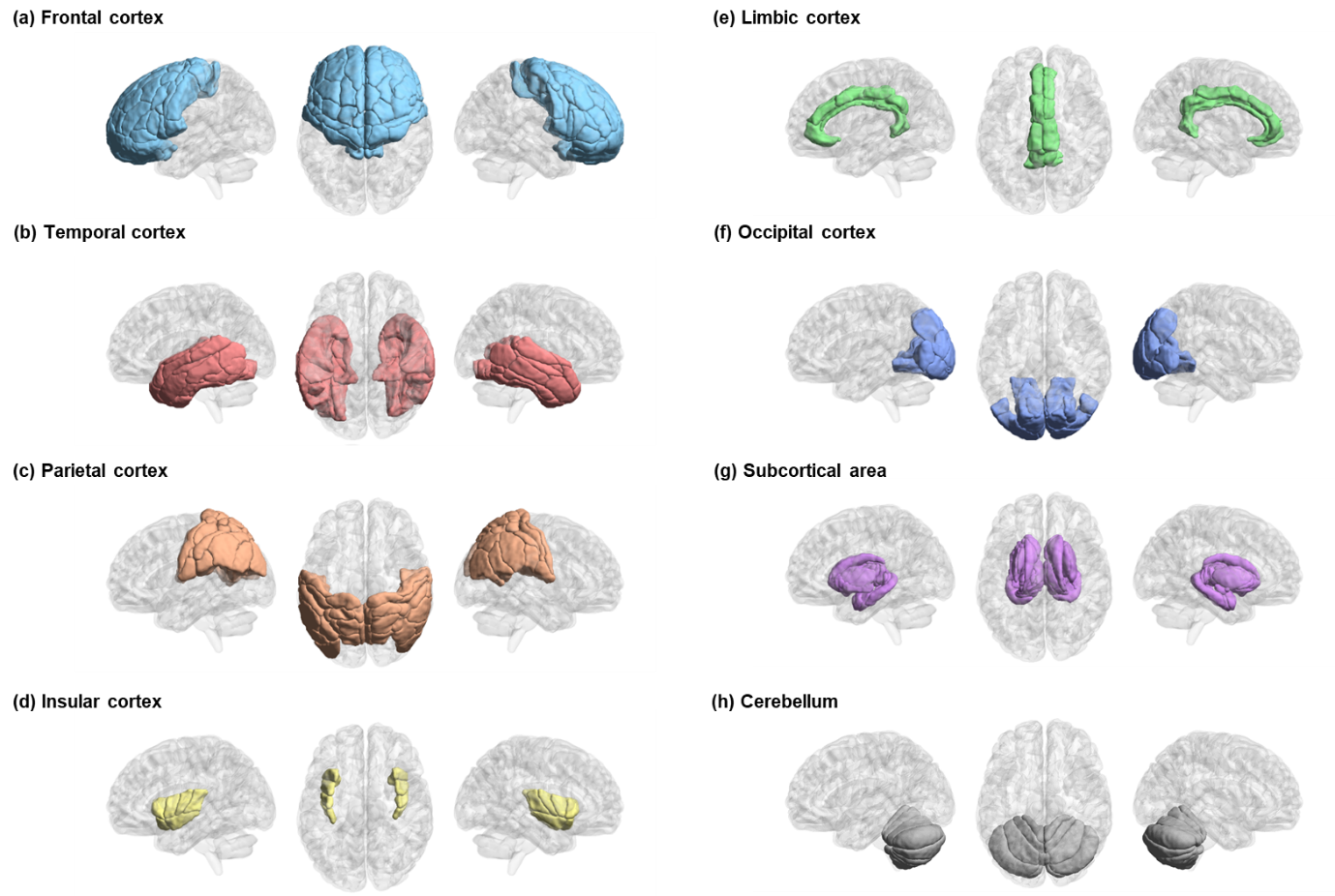

**Supplementary Figure 3. Anatomical labels and their voxels rendered on the 3-dimensional brain.** Eight brain lobes parcellated based on Brainnetome atlas: (a) bilateral frontal lobes, (b) bilateral temporal lobes, (c) bilateral parietal lobes, (d) bilateral insular cortex, (e) bilateral limbic cortex, (f) bilateral occipital lobes, (g) bilateral subcortical area, and (h) cerebellum. Considering the right and the left lobes for the first seven and the cerebellum (containing vermis), fifteen anatomical labels were used for analysis.

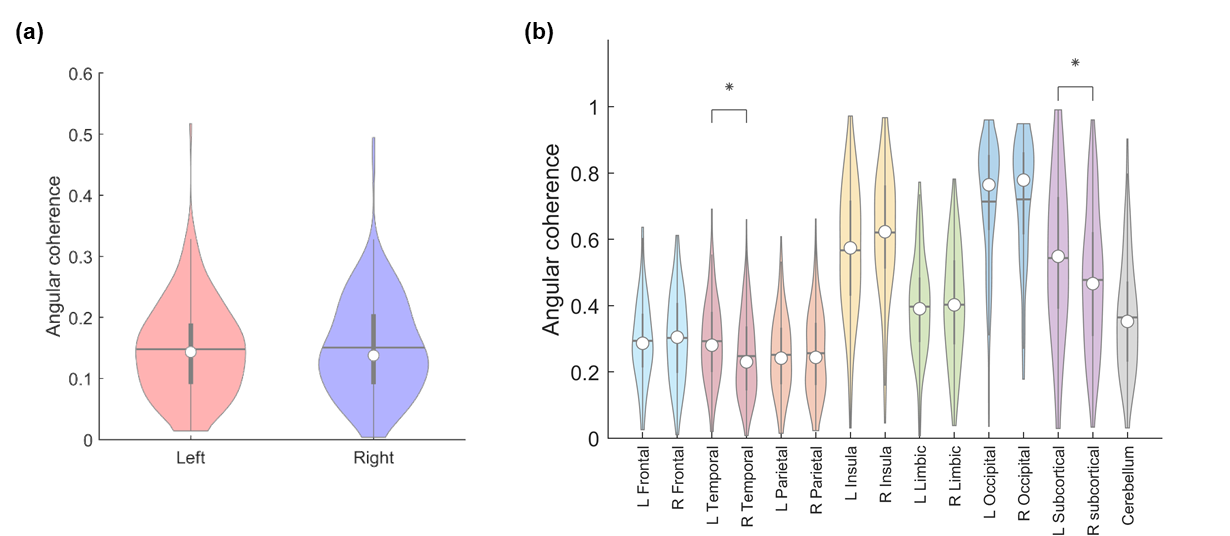

**Supplementary Figure 4. The comparison of angular coherence between the hemispheric-voxels or lobes-voxels of the left and the right hemispheres.** Whether there is any hemispheric laterality in angular coherence was investigated. (a) Designating every voxel into the left or the right hemisphere, there was no significant difference. (b) Differences between the left and the right seven lobes: frontal, temporal, parietal, insula, limbic, occipital lobe, and subcortical region. The temporal lobes and subcortical regions showed significant differences between the left and right (*p* < 0.05, FWER corrected). The left temporal lobe and left subcortical region showed significantly greater angular coherence than the right ones. No difference in angular coherence in the other left and right lobes was found.

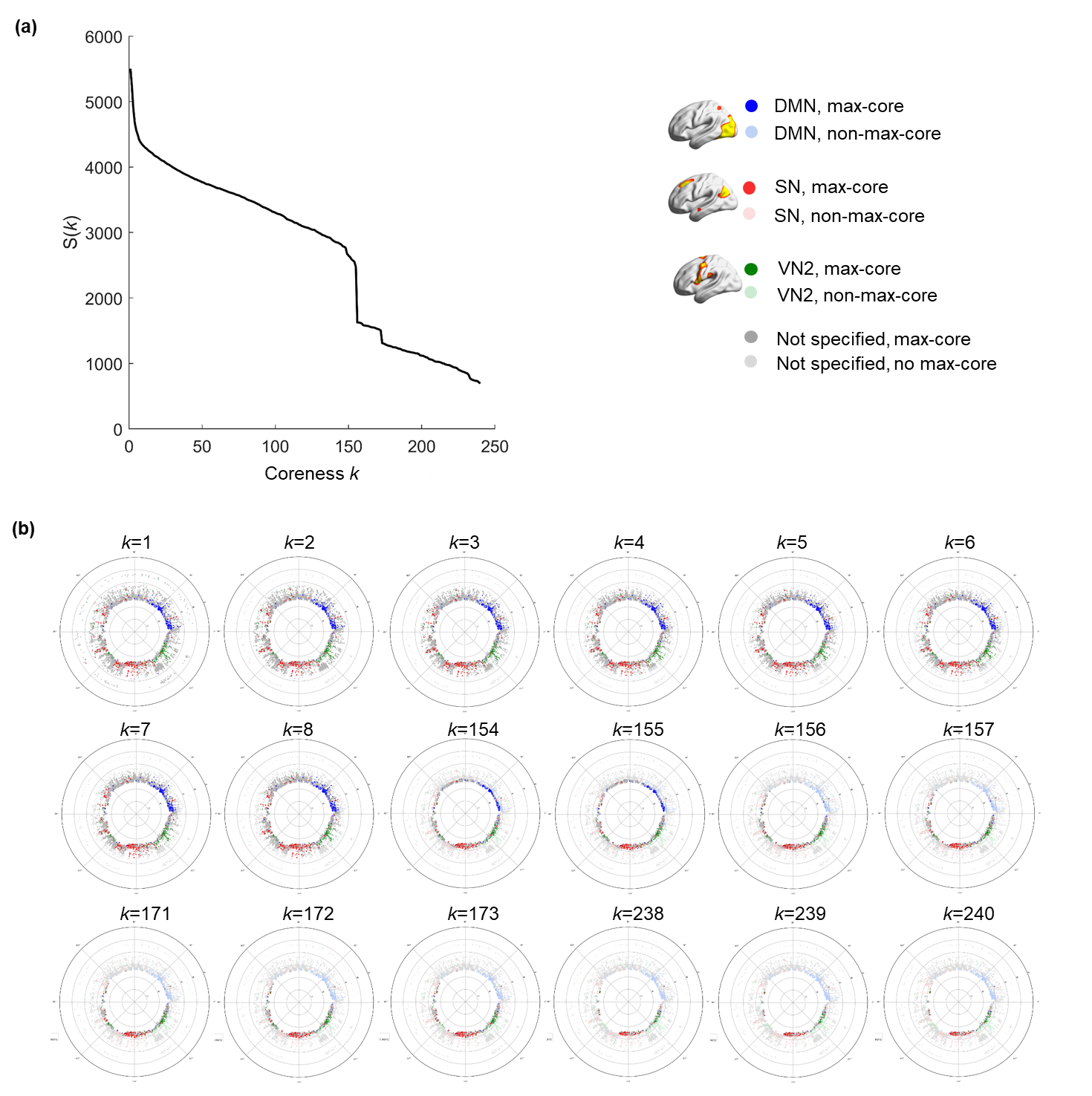

**Supplementary Figure 5. *k*-cores derived from the stepwise *k*-core percolation procedure on the hyperbolically embedded discs.** One can find the changing *k*-core subnetworks of an individual at the sampled *k*-core steps. The *k*-core percolation algorithm removes voxels with *k*-degree, and the algorithm iterates by incrementing coreness *k* by 1. The set of voxels left after each *k*-core percolation step is called *k*-core. The elimination procedure entails recalculating of the degrees of remaining voxels, and the procedure stops when the largest component came to be fragmented into pieces and the largest component can be no more designated. In an example case (subject #100206), (a) The plot shows the size of the *k*-core, S(*k*), according to the coreness *k*. Both gradual and abrupt decreases are shown. (b) The *k*-cores that showed abrupt decreases (*k*=1, 2, 3, 4, 5, 6, 7, 8, 154, 155, 156, 157, 171, 172, 173, 238, 239, 240) were embedded on the hyperbolic discs. Each voxel that belongs to the default mode network (DMN), salience network (SN), and visual network (VN) 2 was colored as blue, red, and green circles, respectively. The gray circles denote voxels that do not belong to DMN, SN, and VN2. The voxel not included in the *k*-cores were shown in pale gray. As *k* increases, more voxels near the edge are eliminated, and an abrupt decrease of S(*k*) involves mass desertion of voxels from then- *k*-cores. Even the voxels with higher degrees near the center were removed nearing the end of *k*-core percolation (examples; *k*=156, 172).

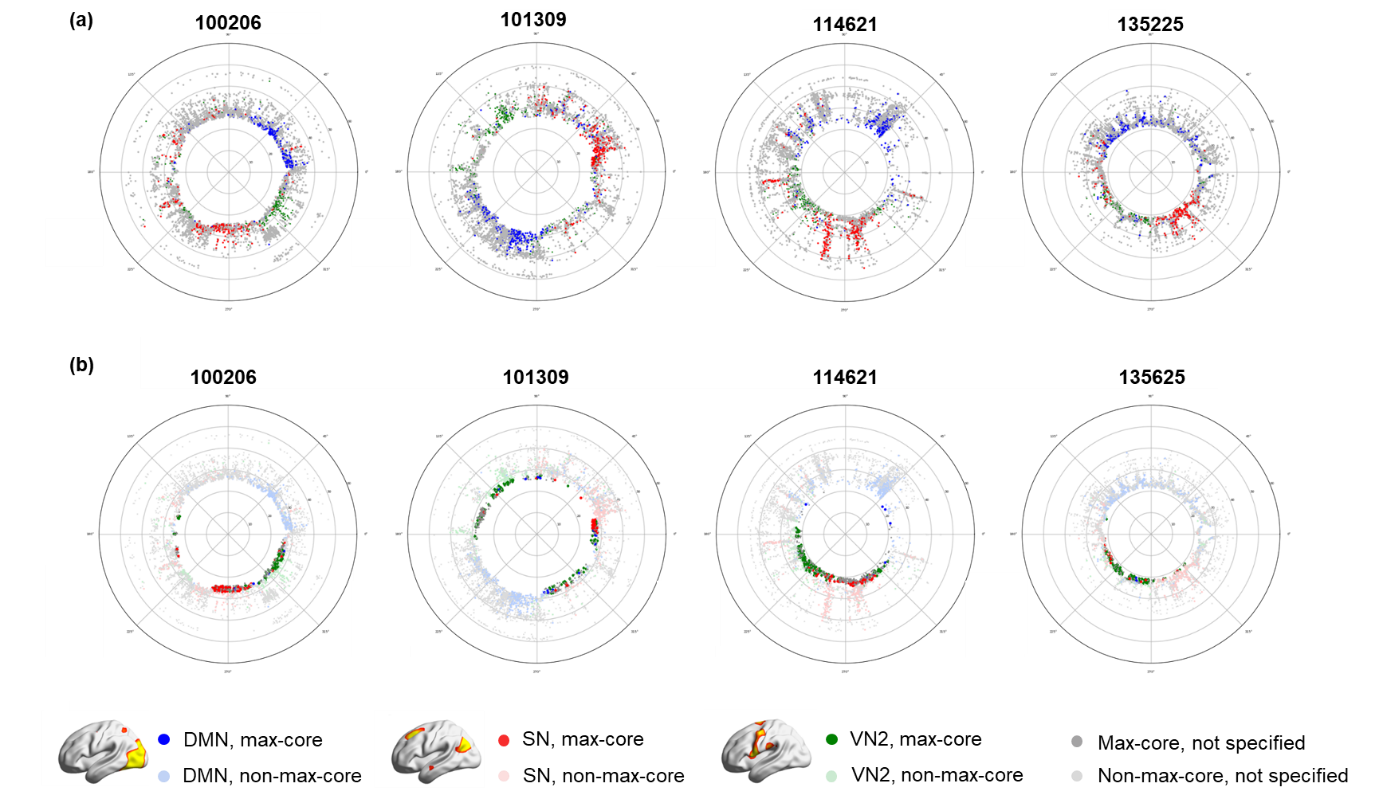

**Supplementary Figure 6. The *k*-cores (*k* = 1 and max) of four individuals** (subject #100206, #101309, #114621, and #135225) **were embedded on their own hyperbolic discs.** (a) Representative voxels on the hyperbolic discs. Voxels that belong to the default mode network (DMN), salience network (SN), and visual network 2 (VN2) are shown as blue, red, and green circles, respectively. The gray circle represents voxels that belong to none of the three. Individuals showed similar but arbitrarily unique patterns of hyperbolic embedding of voxels: larger circle (#100206, #101309), the circle with outer radial pattern (#114621), smaller circle (#135225). Voxels from each network also show various patterns. The SN voxels of an individual (#100206) are distributed in a broad area, whereas those of another individual (#101309) form a cluster. The other (#114621) shows a radial pattern of a cluster consisting of SN voxels. (b) The *k*_max_-core voxels from *k*-core percolation in the above four subjects were visualized on the hyperbolic discs. Voxels that do not belong to *k*_max_-core were shown in a pale color. The numbers of *k*_max_-core voxels and to which independent components they belong vary widely between individuals. Each individual’s ID was shown on top of the disc.

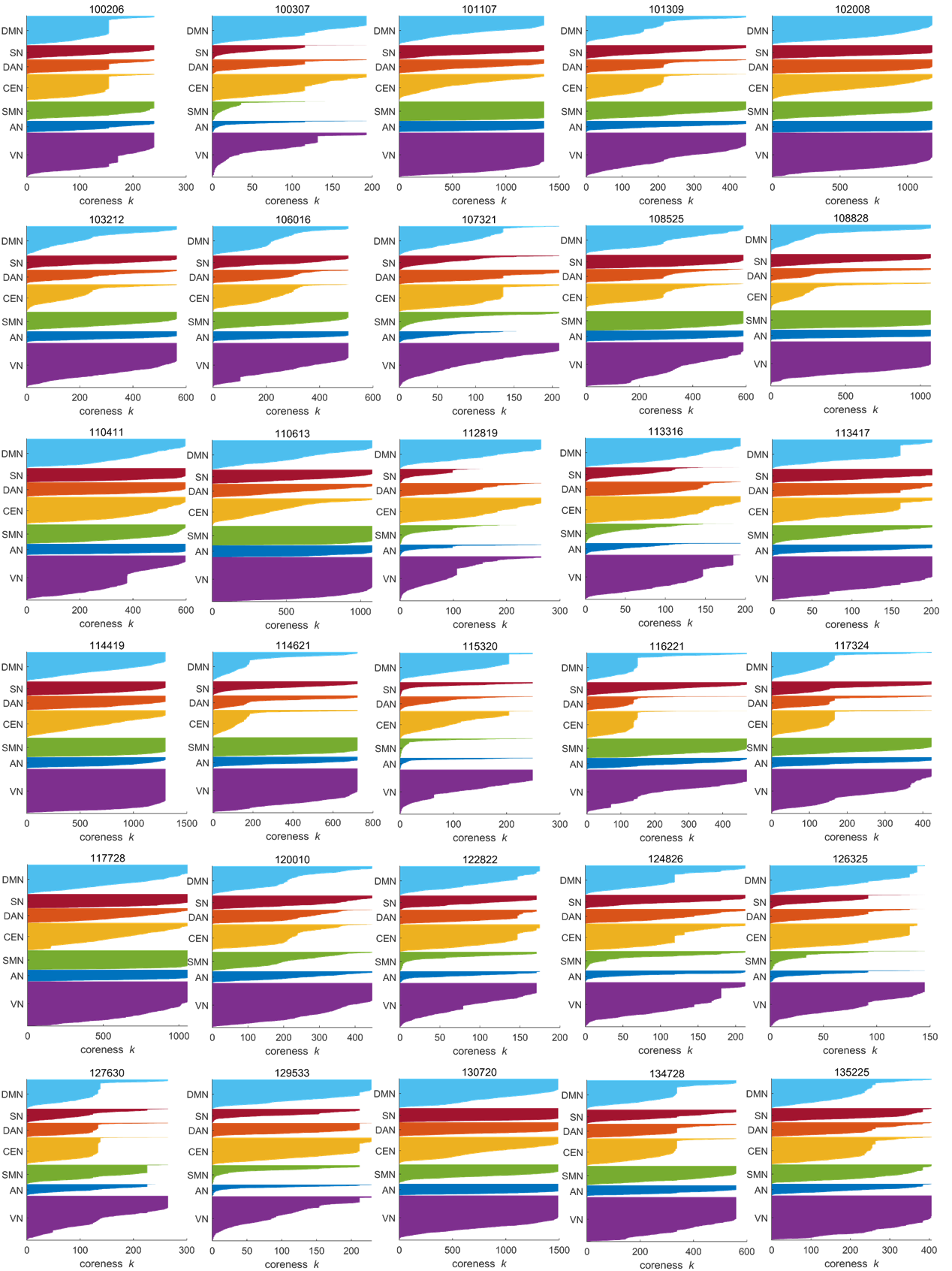

**Supplementary Figure 7. The flags-plots of changes of independent components (ICs)-voxels composition along *k*-core percolation.** The *k*-core percolation peels off the layer of brain network of voxels having degree *k*. As the coreness *k* increases, the size of the *k*-core becomes smaller. The flags-plot shows the change of *k*-cores of each individual according to the coreness *k*. Every voxel that belongs to each IC is shown in the y-axis, and the horizontal bar of each voxel reaches the rightmost until the maximum coreness k. Categorical functional label was used to visualize: default mode network (DMN), salience network (SN), dorsal attention network (DAN), central executive network (CEN), sensorimotor network (SMN), auditory network (AN), and visual network (VN). The voxels of an IC are sorted in descending order of the voxel’s *k* within each flag. Since there are voxels that belong to multiple ICs, the number of voxels in the y-axis is slightly greater than 5,937, the total number of voxels. Every individual shows the unique patterns along *k*-core percolation: 1) abrupt or gradual decrease in the size of *k*-core voxels belonging to each IC along *k*-core percolation, and 2) changes of proportions of voxels that belong to seven categorical ICs.

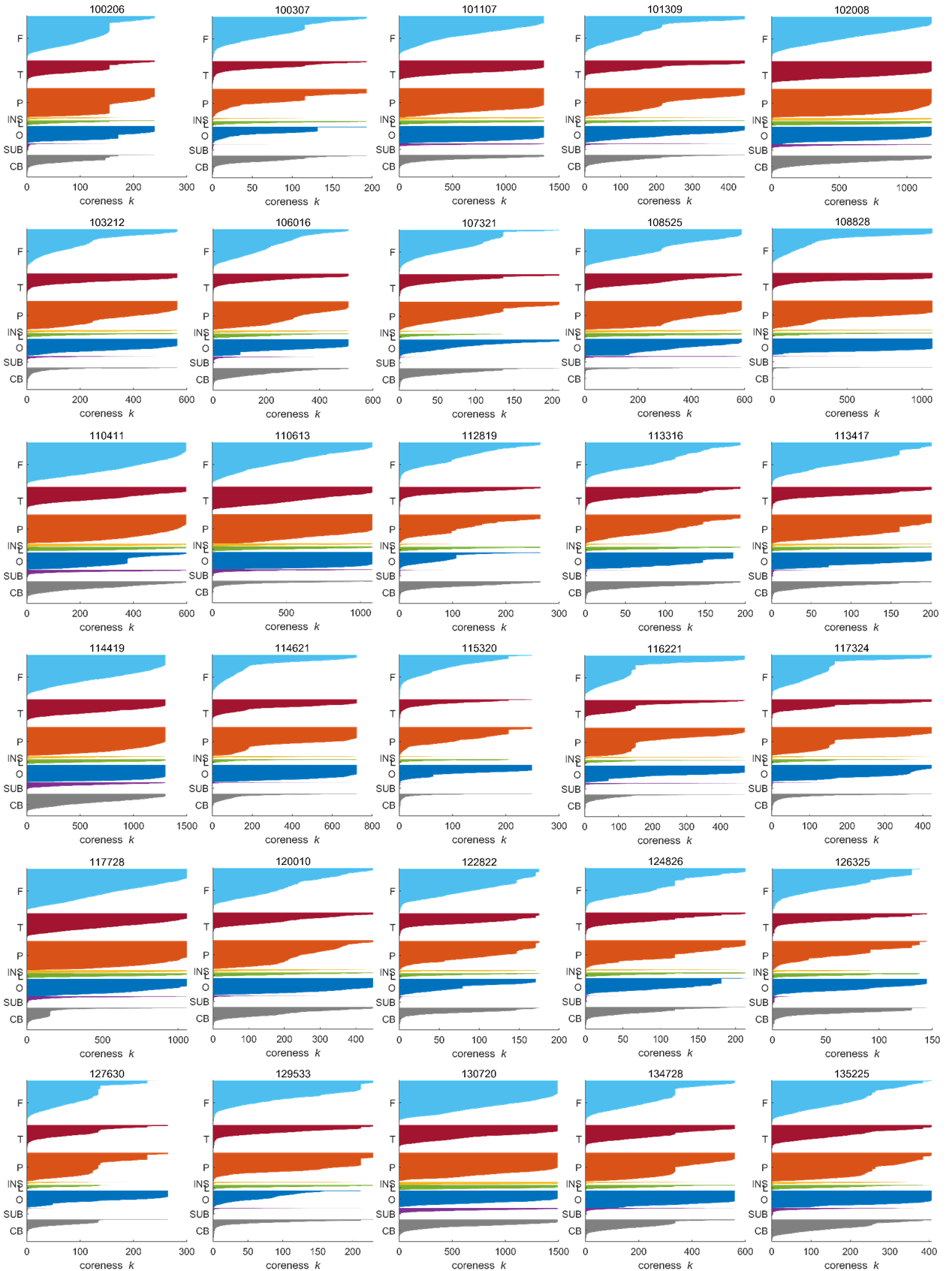

**Supplementary Figure 8. The flags-plots of changes of lobes-voxels composition along *k*-core percolation.** Categorical anatomical labels were used to annotate the voxels into eight lobes: frontal lobe, temporal lobe, parietal lobe, insula, limbic system, occipital lobe, subcortical region, and cerebellum. Each lobe includes voxels from both the left and the right hemispheres. The 5,937 voxels were labeled in the y-axis, and the horizontal bar of each voxel reaches to the rightmost until the maximum coreness *k*. The voxels from each lobe were sorted in descending order for the numbers of lobe-voxels.

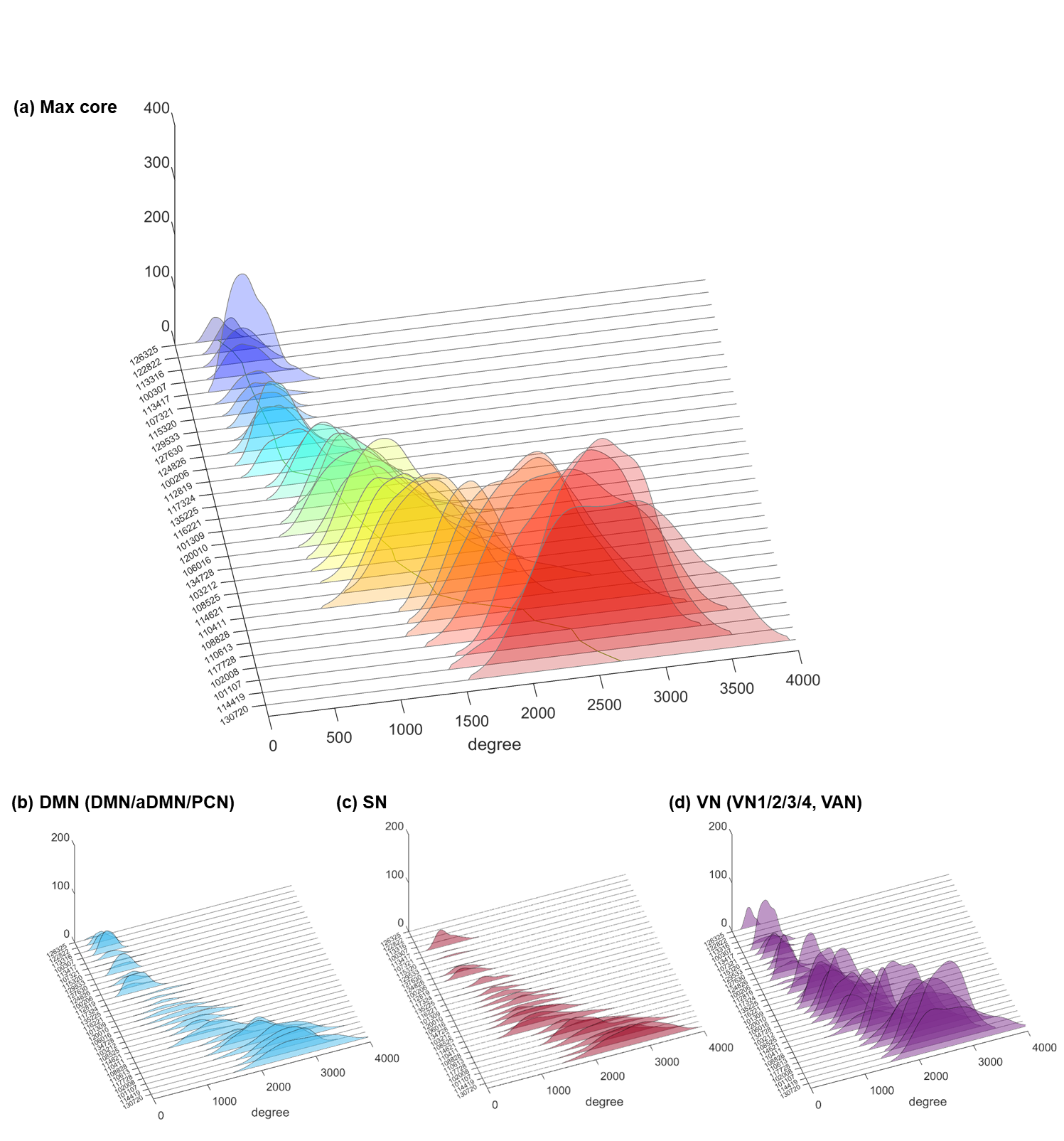

**Supplementary Figure 9. The histogram of *k*_max_-core voxels showing their degree distribution.** The degrees of *k*-core voxels were adopted from their initial adjacency matrix. (a) The histograms of degree distribution were visualized in different colors for individuals. The histograms were sorted in ascending order with the mean degree of the *k*_max_-core voxels. An individual at the top with a deep blue histogram has the lowest voxel degrees, and another with a red at the bottom has the highest. The functional label was used to annotate voxels. (b) The degrees of *k*_max_-core voxels belonging to categorical default mode network (DMN; DMN/anterior DMN (aDMN)/ precuneus network (PCN)), (c) salience network (SN), and (d) visual network (VN; VN1/2/3/4, visual attention network (VAN)) were displayed similarly to (a). The degrees of *k*_max_-core voxels include not only the voxels with dense connections with the colleague *k*_max_-core voxels but also the voxels with lower degrees but effectively connected preferentially with *k*_max_-core connections. In optimal percolation, they called these voxels as influencer nodes.

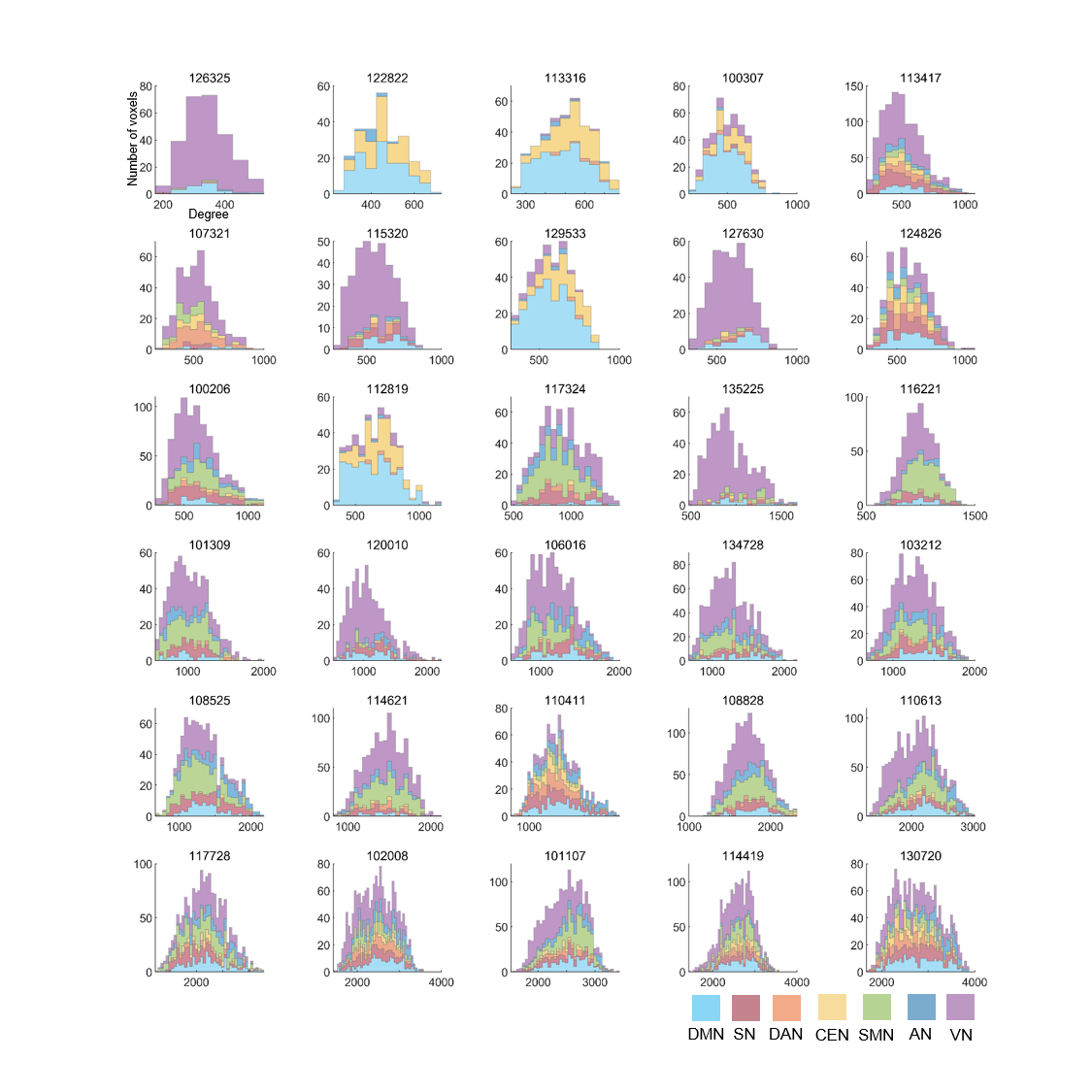

**Supplementary Figure 10. The stacked histogram of the degrees of *k*_max_-core voxels per individual.** The voxel degree was derived from the adjacency matrix, and the affiliation of each voxel was presented in different colors. A *k*_max_-core voxel located on the rightmost side of the histogram refers that the voxel has the greatest degree in the adjacency matrix, indicating that it has many connections with non-*k*_max_-core voxels as well. In contrast, another *k*_max_-core voxel from the left to the rightmost denotes a relatively smaller degree. However, it belongs to *k*_max_-core, implying that it has connections mainly with other *k*_max_-core voxels and only the surplus of degrees are used for connecting itself with non-*k*_max_-core voxels. The histograms of 30 subjects were displayed one by one with sorting in ascending order to the mean degrees of the *k*_max_-core voxels.

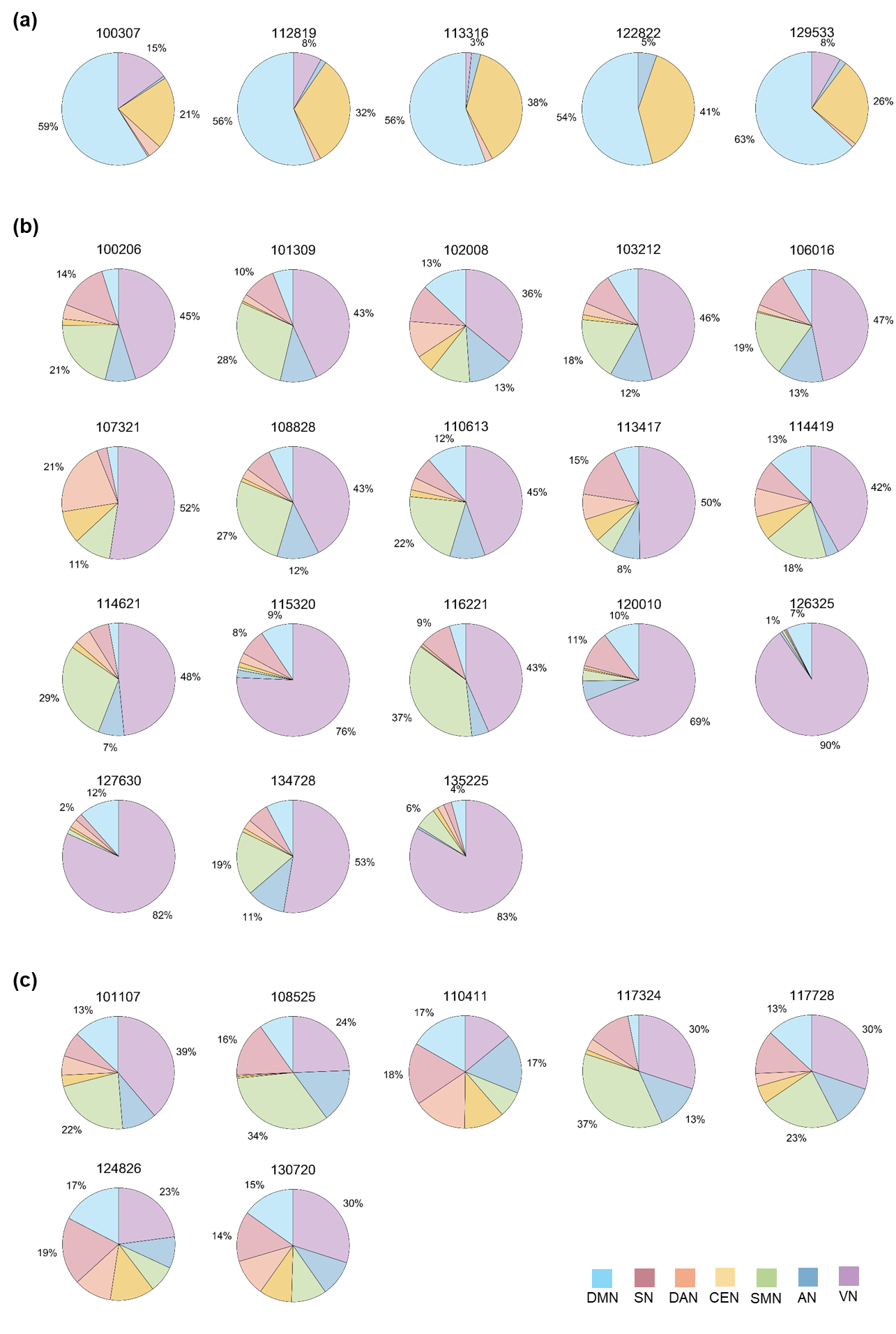

**Supplementary Figure 11. The pie plots showing the ratio of *k*_max_-core voxels of each independent component (IC) over all the *k*_max_-core voxels.** We used a categorical functional label that includes default mode network (DMN), salience network (SN), dorsal attention network (DAN), central executive network (CEN), sensorimotor network (SMN), auditory network (AN), and visual network (VN). The affiliation of *k*_max_-core voxels of each individual was shown in the pie plot, and percentages of the three IC voxels with the greatest sizes were written around the plot. The individuals were assigned into three patterns by the ICs-voxels composition of *k*_max_-core voxels. (a) An individual was grouped as a DMN-dominant if more than 40% of *k*_max_-core voxels belong to DMN. (b) In a VN-dominant, more than 40% of the *k*_max_-core voxels belonged to the VN. (c) A distributed pattern refers that there is no dominant IC for *k*_max_-core voxels. We found five DMN-dominant subjects, 18 VN-dominant subjects, and seven subjects with the distributed pattern.

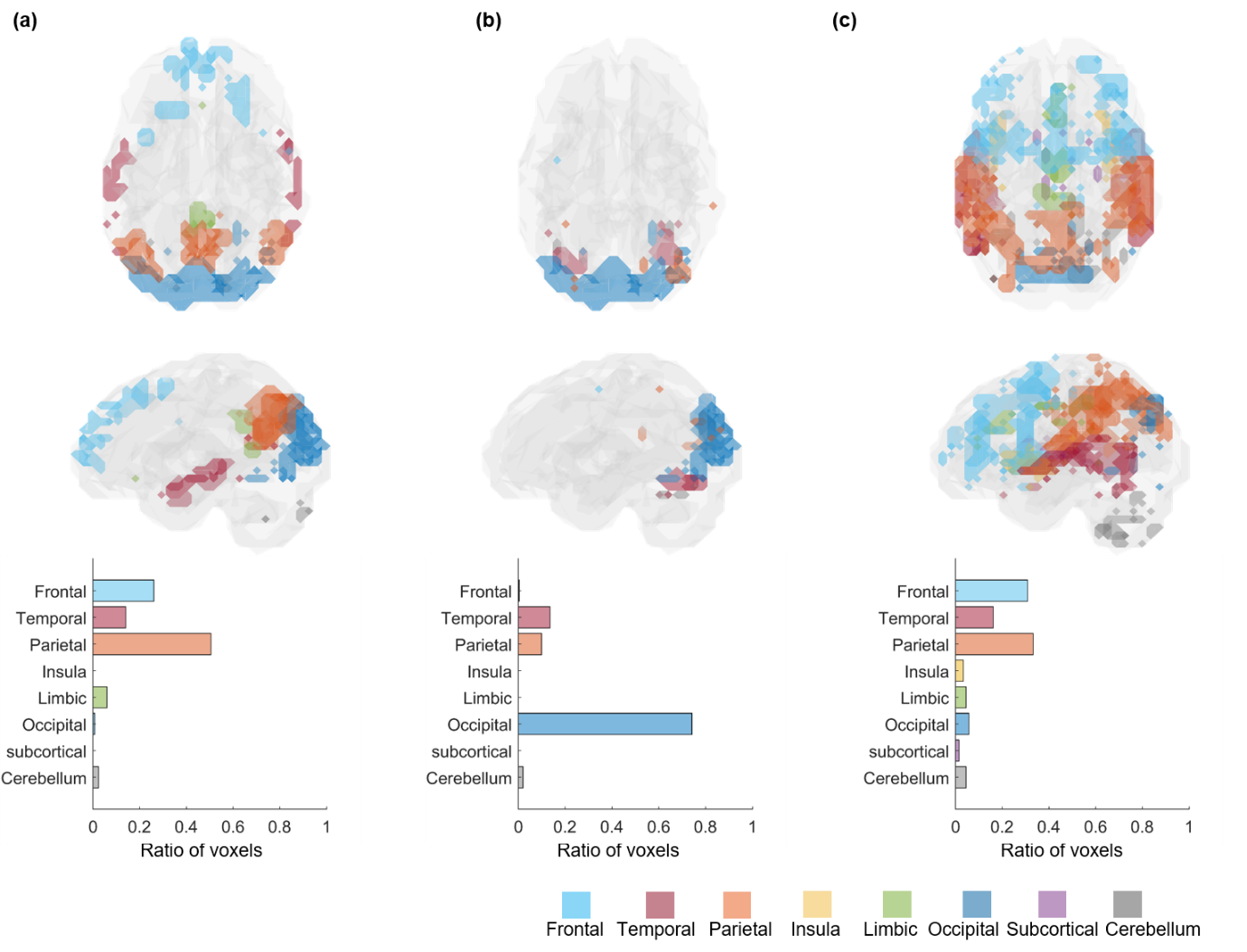

**Supplementary Figure 12. The *k*_max_-core of three individuals from Fig 7 were shown using the anatomical label.** We performed *k*-core percolation to find each individual’s *k*_max_-core voxels and affiliated these *k*_max_-core voxels to the lobes. Categorized anatomical label, including the frontal lobe, temporal lobe, parietal lobe, insula, limbic system, occipital lobe, subcortical region, and cerebellum, was used to annotate *k*_max_-core voxels. Three patterns of *k*_max_-core voxels-ICs composition were found in the three individuals as in Fig 7. (a) In the first individual (129533), *k*_max_-core voxels were mostly in the parietal, temporal, and frontal lobes. (b) In the second individual (126325), *k*_max_-core voxels were mostly in the occipital lobe and a few in the temporal and parietal lobes. (c) In the third individual (110411), *k*_max_-core voxels were distributed in frontal, temporal, and parietal lobes and also in the other lobes. The ratios of the *k*_max_-core voxels of each lobe over the total number were shown as the bar plots at the bottom.

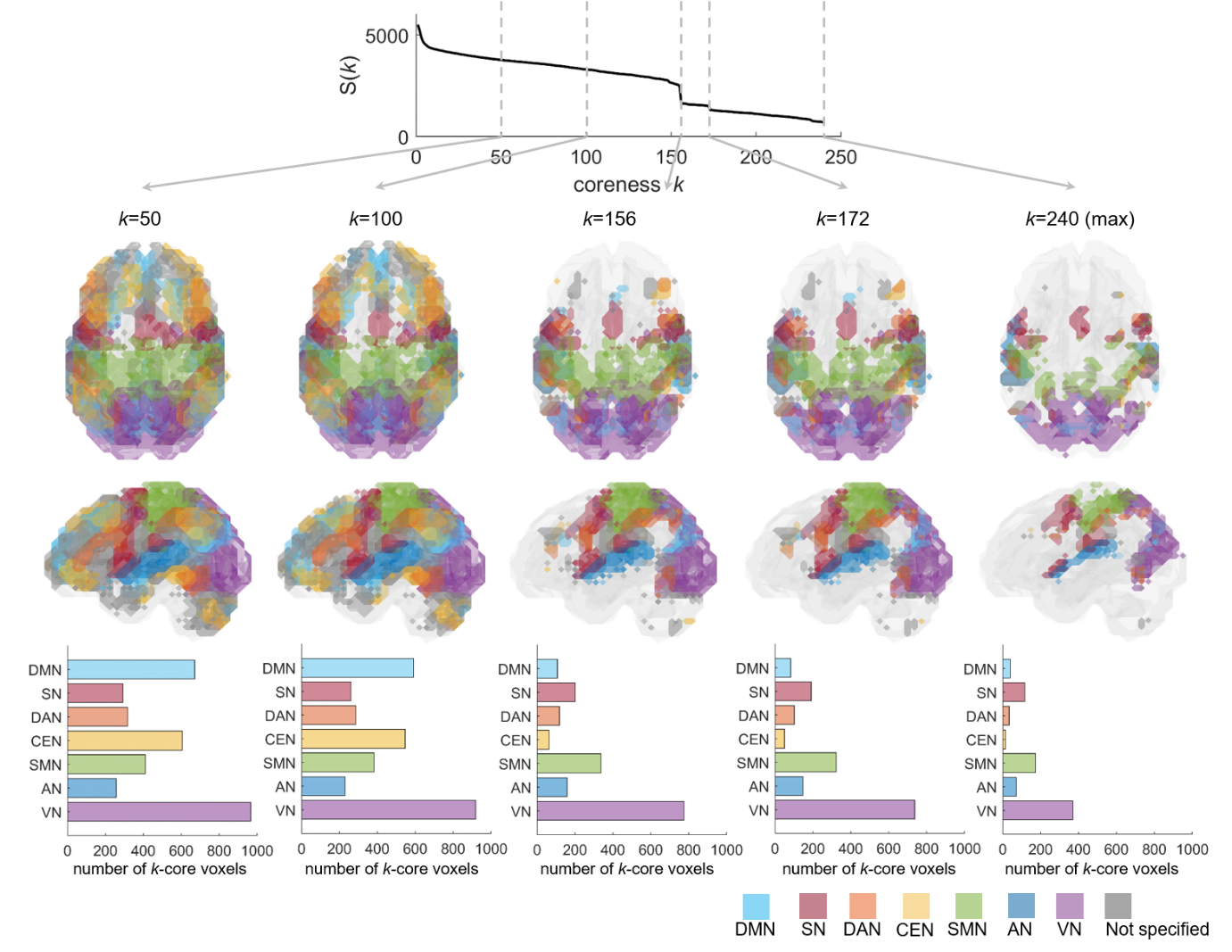

**Supplementary Figure 13. *k*-cores of an individual according to the coreness *k* from *k*-core percolation.** The plot on the top shows the size of *k*-core, S(*k*), according to the coreness *k* while *k*-core percolation proceeds of this individual (100206). We used a categorical functional label, which includes seven combined independent components (ICs), to classify *k*_max_-core voxels. Each voxel is printed on the 3-dimensional brain in corresponding colors in the middle. The bar plots on the bottom show the number of *k*_max_-core voxels that belong to each IC. As coreness *k* increases, the number of voxels belonging to each IC decreases, and at the step maximum, there remain *k*_max_-core voxels. Voxels belonging to the visual network (VN) are more than 900 when *k* is 1 and comes to be less than 400 at the *k*_max_ step. However, VN yet occupies the majority since the number of *k*_max_-core voxels belonging to other ICs comes to be far fewer.
